# Anticorrelated inter-network electrophysiological activity varies dynamically with attentional performance and behavioral states

**DOI:** 10.1101/503193

**Authors:** Aaron Kucyi, Amy Daitch, Omri Raccah, Baotian Zhao, Chao Zhang, Michael Esterman, Michael Zeineh, Casey H. Halpern, Kai Zhang, Jianguo Zhang, Josef Parvizi

## Abstract

The default mode network (DMN) is thought to exhibit infraslow anticorrelated activity with dorsal attention (DAN) and salience (SN) networks across various behavioral states. To investigate the dynamics of activity across these networks on a finer timescale, we used human intracranial electroencephalography with simultaneous recordings within core nodes of the three networks. During attentional task performance, the three sites showed dissociable profiles of high-frequency broadband activity. Anticorrelated infraslow fluctuations of this activity were found across networks during task performance but also intermittently emerged during rest and sleep in concert with the expression of task-like network-level topographic patterns. Critically, on a finer timescale, DAN and SN activations preceded DMN deactivations by hundreds of milliseconds. Moreover, greater lagged, but not zero-lag, anticorrelation between DAN and DMN activity was associated with better attentional performance. These findings have implications for interpreting antagonistic network relationships and confirm the behavioral importance of time-lagged inter-network interactions.

## Introduction

The brain structures important for attention have long been described as a set of discrete components of networks, each of which serves a unique role in cognitive interactions with the sensory environment (Posner and Petersen, 1990). The *dorsal attention network* (DAN) is implicated in goal-directed (top-down) attention (Corbetta and Shulman, 2002), and the *salience network* (SN) is implicated in stimulus-driven (bottom-up) attention and cognitive control (Seeley et al., 2007, Uddin, 2015). Given functional neuroimaging evidence, these described networks exhibit opposing activity with yet another set of regions that constitute the *default mode network* (DMN) (Raichle et al., 2001, Fox et al., 2005), which unlike DAN and SN structures, tends to show deactivation during conditions involving externally-oriented attention (Buckner et al., 2008). It has been hypothesized that anticorrelated activity between the DMN and DAN/SN reflects a competition for control over shared computational resources and could be a marker of one’s level of engagement with externally-oriented tasks (Sonuga-Barke and Castellanos, 2007, Anticevic et al., 2012).

In addition to task-dependent activity, fMRI studies during wakeful rest have shown spontaneous anticorrelated DMN-DAN/SN activity in infraslow (<0.1 Hz) fluctuations of blood-oxygen-level-dependent (BOLD) signals (Fox et al., 2005, Fransson, 2005) – a finding that has remained contentious (Murphy and Fox, 2017). Persistence of such anticorrelated activity in task-free states would potentially suggest that functionally competing systems, characterized by continual switching between internally- and externally-biased modes of attention, are an intrinsic property of the brain (Buckner et al., 2013, Honey et al., 2018).

To date, however, evidence for anticorrelated activity has relied almost exclusively on fMRI, which offers limited temporal resolution and necessitates data preprocessing that may bias estimates of negative correlations (Chang and Glover, 2009, Murphy et al., 2009). Intracranial electroencephalography (iEEG) in human subjects offers anatomical precision, high temporal resolution, and sensitivity to activity in the high-frequency broadband (HFB, also known as high gamma) range (~70-170 Hz)– a well-established correlate of the BOLD signal and neuronal population spiking (Parvizi and Kastner, 2018). A handful of iEEG studies involving recordings from putative DMN and DAN nodes have shown task-evoked HFB responses that resemble the antagonistic inter-network patterns observed in fMRI (Ossandon et al., 2011, Ramot et al., 2012, Raccah et al., 2018). In addition, resting state iEEG has revealed correlates of the DMN and DAN (Foster et al., 2015, Hacker et al., 2017, Kucyi et al., 2018a) and that a subset of region pairs with resting BOLD anticorrelations exhibit weaker, but significant anticorrelations of slow (0.1-1 Hz) HFB activity (Keller et al., 2013).

The extant iEEG evidence provides promising initial electrophysiological validation of antagonistic inter-network interactions, but beyond confirming fMRI findings, critical open questions remain: Is infra-slow anticorrelated activity between task-responsive DMN, DAN and SN neuronal populations found in task-free states and in specific frequency components of electrophysiological signals? Do nodes of the DMN, DAN and SN exhibit distinguishable, systematic temporal profiles of task-evoked electrophysiological activity? Are time-resolved antagonistic interactions relevant to intra-individual variations in attentional task performance?

Here we report an iEEG investigation of anticorrelated brain networks across multiple sessions of attentional task performance, wakeful rest and sleep. We localized iEEG recording sites in key nodes of the DMN, DAN, and SN through a rigorous survey of anatomical boundaries, iEEG response profiles, and within-individual resting-state fMRI connectivity. We then examined how inter-network iEEG interactions vary as a function of attentional (task performance) and behavioral (active-task versus rest versus sleep) states. *We hypothesized that temporal features of electrophysiological anticorrelated activity would be associated with fluctuations in sustained attention and that the overall magnitude of anticorrelated activity would co-vary within and across behavioral states despite a constrained wider network-level spatial organization*.

## Results

### Unique iEEG cohort

We obtained iEEG recordings from a total of 896 recording sites in seven participants (S1-S7, five with depth electrode and two with subdural recordings) undergoing treatment for focal epilepsy. Data reported here are from regions void of pathological activity and outside each participant’s seizure zone. An average of 78±14% of channels per participant were retained for analysis, a percentage that is within the typical range (Parvizi and Kastner, 2018). Each subject had simultaneous electrode coverage across key nodes of DMN, DAN and SN, as defined based on anatomical boundaries and, when available (n=5), confirmed with individual-level resting-state fMRI (see Methods). Specifically, we focused on recordings from three regions: 1) posteromedial cortex (PMC) within the DMN (Raichle et al., 2001); 2) dorsal posterior parietal cortex (dPPC), including superior parietal lobule and intraparietal sulcus, within the DAN (Corbetta and Shulman, 2002); and 3) dorsal anterior insular cortex (dAIC) within the SN (Seeley et al., 2007). Our cohort was unique and carefully selected from a larger group of patients based on the presence of recording sites within the three regions of interest. Implantation of intracranial electrodes in all subjects were solely based on clinical needs, and the location and number of electrodes implanted in each case were decided by a collective agreement of medical staff at Stanford Medical Center (6 subjects) and Beijing Tiantan Hospital, Capital Medical University (1 subject).

Subjects performed between four to eight sessions (total duration range: 24-48 minutes per subject) of the *Gradual-onset Continuous Performance Task* (GradCPT), a test of sustained attention that has reliably been associated with anticorrelated DMN versus DAN/SN activity in fMRI studies (Esterman et al., 2013, Kucyi et al., 2016, Fortenbaugh et al., 2018). Gradually changing images of scenes were presented every 800 ms, and subjects were instructed to respond with a button press to city (frequent) but not to mountain (infrequent) scenes (**Figure 1A**). The task requires sustained attention and withholding of a habitual response to infrequent target events. We defined behavioral performance within each session of the experiment by the measure of sensitivity (*d*′) which is based on the accuracy of task performance (accounting for both hits and false alarms) (Fortenbaugh et al., 2015, Rosenberg et al., 2016). We found that performance varied from session to session within subjects (**Table S1**). In three subjects, additional recordings were obtained during wakeful rest and sleep sessions of similar durations to those obtained during task performance.

**Figure 1.**
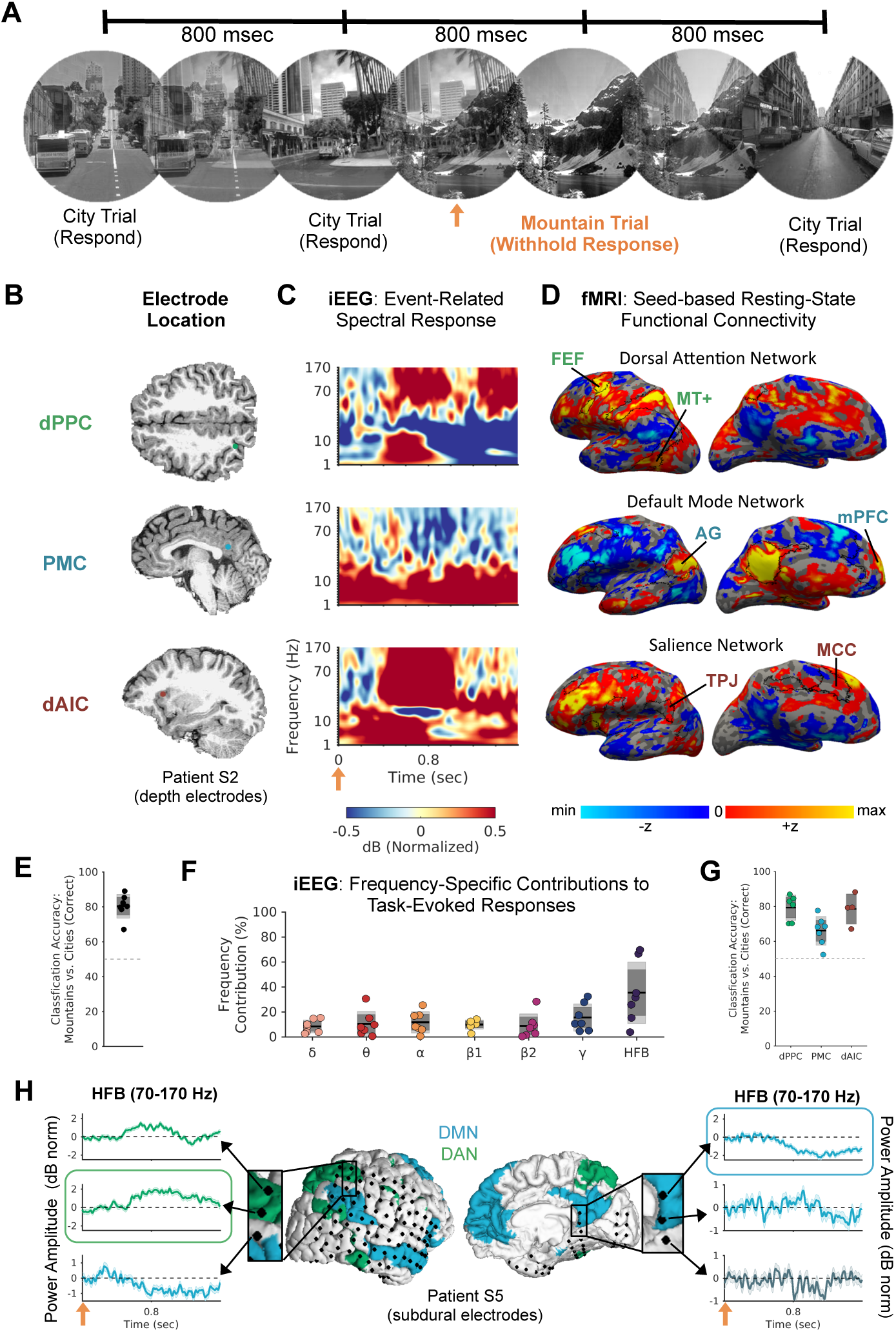
Task paradigm and functional localization of electrode sites in three regions of interest. **A)** The Gradual-Onset Continuous Performance Task. City and mountain scene images faded continuously from one image to the next every 800 msec. Subjects were instructed to press a button when they saw cities but to withhold responses to (rare) mountains. Trial onset (orange arrow) was the time at which stimulus fade-in was initiated. **B)** Anatomical locations of electrode contacts implanted in three regions of interest that showed peak-responsive high-frequency broadband (HFB; 70–170 Hz) responses during withheld responses to mountains (correct omissions) in an example subject. **C)** Time-frequency plots for regions shown in B), highlighting spectral changes during correct omissions. In the dPPC (top) and dAIC (bottom), HFB increases were found, whereas in the PMC (middle), an HFB decrease was found. **D)** Resting-state functional connectivity (based on pre-operative fMRI) from seed locations at the dPPC, PMC and dAIC electrode locations shown in B). Red/yellow indicates positively correlated regions; blue/light blue regions indicates negatively correlated regions (z scores based a general linear model analysis, thresholded arbitrarily for display purposes). Black outlines on the cortical surface indicate the boundaries of the dorsal attention network (top), default mode network (middle), and salience network (bottom) based on the Yeo et al. (2011) population-level atlas registered to the subject’s brain. **E)** Classification accuracy (correct omissions versus correct commissions) in all seven subjects based on multikernel learning analysis (full model with seven frequency ranges and all channels within dPPC, dAIC, and PMC). Frequency ranges correspond to δ (1–3 Hz), θ (4–7 Hz), α (8–12 Hz), β1 (13–29 Hz), β2 (30–39 Hz), γ (40–70 Hz) and HFB (70–170 Hz). **F)** Contributions of the power amplitudes of distinct frequency bands to the classification accuracy shown in E). **G)** Classification accuracy, as in E), but for models that include channels only within dPPC, dAIC or PMC. **H)** Subdural electrodes plotted on the cortical surface with an overlay of the Yeo atlas’ DMN (blue) and DAN (green) in patient S5. Time series plots illustrate how neighboring channels may display diverging HFB response profiles that partially correspond to network identity (peak-responsive channels in the dPPC and PMC, respectively, are outlined in green and blue). See also **Figure S1** and **Figure S2**.

### Functional localization of electrophysiological antagonistic networks

We functionally localized electrode contacts of interest within the PMC, dPPC and dAIC anatomical boundaries (**Figure 1B**, see Methods for details). These sites were chosen as core nodes of three networks of interest, namely the DMN, DAN, and SN respectively. We analyzed electrophysiological activity during withheld presses to infrequent targets (correct omissions) relative to presses for frequent non-targets (correct commissions) (**Figure 1C**).

In our analysis, we adopted a recently described data-driven multivariate approach to decompose the relative contributions of distinct bands of the iEEG signal to task-evoked responses (Schrouff et al., 2016). Specifically, for correct omission (mountain) and correct commission (city) trials, we generated kernels of similarity across trials for each channel that was anatomically within at least one of the ROIs (dPPC, PMC, dAIC) and the power amplitudes of seven frequency bands [*δ* (1-3 Hz), *θ* (4-7 Hz), *α* (8-12 Hz), *β1* (13-29 Hz), *β2* (30-39 Hz), *γ* (40-70 Hz) HFB (70-170 Hz)]. Using these features (i.e., 7 frequencies x number of ROI channels in a given subject), multikernel learning analyses revealed significant classification accuracy within each subject (*M*±*SD*=80.7±7.0%, p<0.01 in all cases) (**Figure 1E**). Across frequencies, HFB features had the highest mean contributions to classification accuracy across subjects, although lower frequencies also contributed (**Figure 1F**). Multikernel learning models based on single ROIs (and the seven frequencies) revealed that classification accuracy was strongest for dPPC (*M*=79.3%) and dAIC (*M*=78.5%) and weakest for PMC (*M*=66.1%) activity (**Figure 1G**). Based on these findings, we focus our central further iEEG analyses on HFB activity but also consider other frequency ranges.

As noted, HFB (~70-170 Hz) is a well-established correlate of the BOLD signal (and neuronal population spiking) (Parvizi and Kastner, 2018). Based on previous fMRI findings (Fortenbaugh et al., 2018), we expected decreased HFB activity in the PMC following targets relative to non-targets, while in dPPC and dAIC, we expected greater HFB activity following targets relative to non-targets. In each subject (n=7), we successfully identified peak-responsive channels within the PMC that showed temporal clusters of significantly decreased HFB power following target compared to non-target onsets (Monte Carlo *p*<0.002 in all instances, corrected for number of channels within ROI using cluster-based permutation testing). In each participant with dPPC (n=6) and dAIC (n=4) coverage, we also identified temporal clusters of significantly increased HFB power within the dPPC (Monte Carlo *p*<0.001 in all instances) and dAIC (Monte Carlo *p*<0.001 in all instances) for target compared to non-target trials (see **Figure S1A** for all subjects). In the peak-responsive channels, spectrograms suggested that deactivations (PMC) and activations (dPPC, dAIC) were consistent within the HFB range, although responses were also evident in lower frequency ranges (See **Figure S1B** for all subjects).

In addition to demonstrating task-evoked iEEG response profiles, we confirmed the network membership of peak-responsive channels within each ROI using resting-state fMRI. To do so, we extracted the BOLD time series from the channels’ locations and performed seed-based functional connectivity analysis (Biswal et al., 1995) to identify remote brain regions with correlated infraslow BOLD activity. In the five subjects who underwent resting-state fMRI, well-known features of the DMN, DAN and SN were found at the individual level (single-subject example in **Figure 1D**; see **Figure S1C** for all subjects). Specifically, for the PMC peak-deactive iEEG sites, selectively correlated BOLD activity was found with established nodes of the DMN including medial prefrontal cortex and angular gyrus. For the dPPC peak-active iEEG sites, correlated BOLD activity was found with nodes of the DAN including frontal eye fields and area MT+. For dAIC peak-active iEEG sites, correlated BOLD activity was found with the mid-cingulate cortex and anterior temporoparietal junction within SN. Also apparent in these BOLD network maps were DMN-DAN and DMN-SN anticorrelations, which were obtained using global signal regression but which also were apparent when using an alternative preprocessing pipeline (**Figure S2**).

Overlays of DMN, DAN and SN templates, derived from a population-level atlas (Yeo et al., 2011), showed concordance between BOLD networks in each individual subject and those found in neurotypical adults (black outlines on cortical surfaces in **Figures 1D and S1C**). In the two subjects that did not undergo fMRI, registration of the Yeo atlas to individual cortical surfaces suggested that peak-responsive PMC, dPPC, and dAIC channels were respectively within the boundaries of the fMRI-defined DMN, DAN and SN. The correspondence between iEEG response profile and fMRI network identity could be illustrated in cases where subdural electrodes densely covered areas near network boundaries. For example, in S5, dPPC and PMC responsive sites were found within the Yeo atlas’ DAN and DMN, respectively, but HFB responses at adjacent sites a few millimeters away were absent or distinct (**Figure 1H**). These network-wide spatial profiles, together with the observed task-evoked iEEG response profiles, suggest that the general functions and connectivity of the PMC, dPPC and dAIC were preserved within participants.

### Dissociable DAN and SN response profiles

As noted, both dPPC and dAIC exhibited similarly increased electrophysiological responses in the HFB range during correct omission trials (**Figure S3A**). However, during commission errors that signify lapses of attention (i.e., when participants incorrectly pressed the key button to infrequent mountain targets), we observed a clear divergence between DAN and SN sites. As expected from fMRI studies (Ham et al., 2013, Neta et al., 2015, Fortenbaugh et al., 2018), during commission errors compared with correct omission trials, the dAIC showed temporal clusters of significantly greater HFB activation in *all* 4 subjects with dAIC coverage (Monte Carlo *p*<0.05 in all cases) whereas the dPPC showed no significant differences in any of the 6 subjects with dPPC coverage (**Figure 2A and 2C**). The error-related HFB increases in the dAIC appeared later and more sustained compared with those found for correct omission trials (**Figure 2C and 2D**). The PMC showed largely similar HFB deactivation for correct omission and commission error trials, although in 2 out of 7 PMC subjects, temporal clusters of greater deactivation for correct trials were found (Monte Carlo *p*<0.05 in both cases) (**Figure 2B**). Taken together, these findings further confirm that dPPC and dAIC recording sites were within functionally dissociable networks.

**Figure 2.**
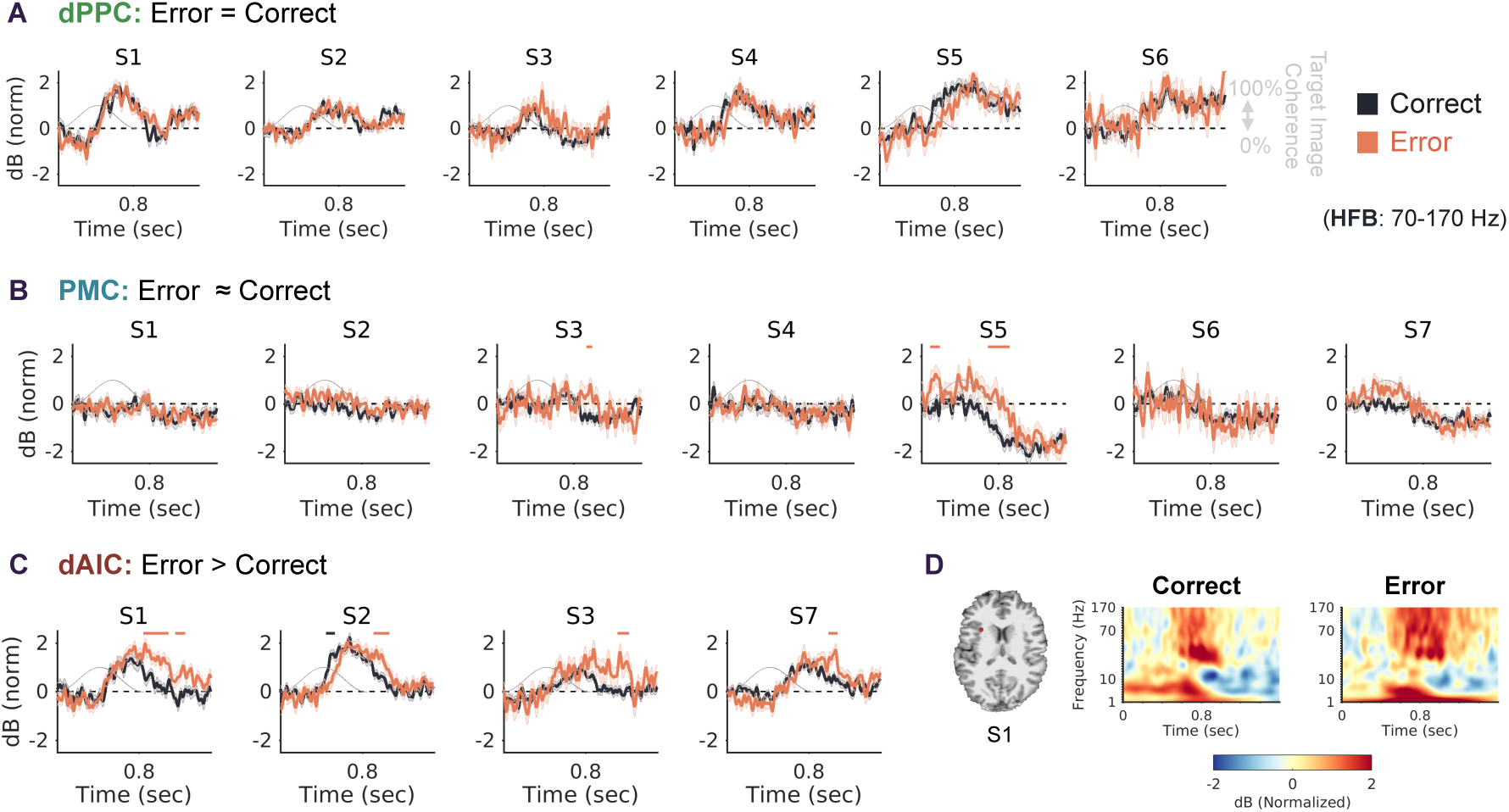
High-frequency broadband (70-170Hz) electrophysiological responses during correct omissions (red) and commission errors (black) in all subjects. **A)** Comparisons for peak-responsive (HFB increase) channels in dorsal posterior parietal cortex (dPPC). No significant differences between correct and errors trials were found in all 6 dPPC subjects. The gray sine wave indicates the target image coherence (mountain fading in to full coherence at t=400 msec, and then fading back out) during the trial. **B)** Comparisons for peak-responsive (HFB decrease) channels in the posteromedial cortex (PMC). In 2 out of *7* PMC subjects, temporal clusters were found of significantly greater HFB power during correct compared to error trials (red horizontal lines above plots; Monte Carlo p<0.05). **C)** Comparisons for peak-responsive (HFB increase) channels in the dorsal anterior insular cortex (dAIC). In all 4 dAIC subjects, temporal clusters were found of significantly greater HFB power during correct compared to error trials (red horizontal lines above plots; Monte Carlo p<0.05). In 1 out of 4 subjects, a temporal cluster was found of greater HFB power during error compared to correct trials (black horizontal line above plots; Monte Carlo p<0.05). **D)** Location of peak-responsive dAIC electrode and spectrograms for correct and error trials in an example subject (S1). For error compared to correct trials, a stronger and more sustained increase in power amplitude can be seen in the HFB range.

### Diminished anticorrelated iEEG activity during rest and sleep states

Past fMRI studies have suggested the possibility of anti-correlated inter-network activity during non-task states (Fox et al., 2005, Fransson, 2005) – but these findings have remained disputed (Murphy and Fox, 2017). Since task-independent anticorrelated inter-network activity in fMRI has relied on infraslow (<0.1 Hz) fluctuations of BOLD signals, we used comparable parameters to explore the presence of task-independent intrinsic anticorrelations between PMC and dPPC and between PMC and dAIC. For this, we selected the same functionally localized (peak-responsive) PMC, dPPC and dAIC sites shown in the previous analyses and assessed whether infraslow (<0.1 Hz) HFB envelope anticorrelation was found between them.

In three patients (S1, S2, S6) with rest and sleep (**Figure S4**) recordings of similar total duration to task recordings (24-48 minutes per state; **Tables S2, S3 and S4**), we split recordings into independent 100-second windows and compared infraslow HFB functional connectivity across states. We found that within each subject, there was a significant interaction between behavioral state and infraslow HFB functional connectivity for dPPC-PMC (*F*>10, *p*<0.001 in all cases) and dAIC-PMC (*F*>10, *p*<0.001 in all cases) channel pairs. Though some time windows with anticorrelated activity were found during rest and sleep, consistent anticorrelation across windows was reliably detected only in the task state for both dPPC-PMC (**Figure 3A**) and dAIC-PMC (**Figure 3C**) channel pairs. In all subjects, infraslow HFB functional connectivity in task was significantly lower than that in rest for dPPC-PMC (*p*<0.005 in all cases) and dAIC-PMC (*p*<0.005 in all cases) channel pairs. Infraslow HFB functional connectivity was also significantly lower in task than in sleep for dPPC-PMC (*p*<0.001 in all cases) and dAIC-PMC (*p*<0.001 in all cases) channel pairs. Differences between rest and sleep were found less consistently for both dPPC-PMC (S1: *p*=0.03; S2: *p*=0.14; S6: *p*=0.81) and dAIC-PMC (S1: *p*=0.89; S2: *p*=0.046) channel pairs.

**Figure 3.**
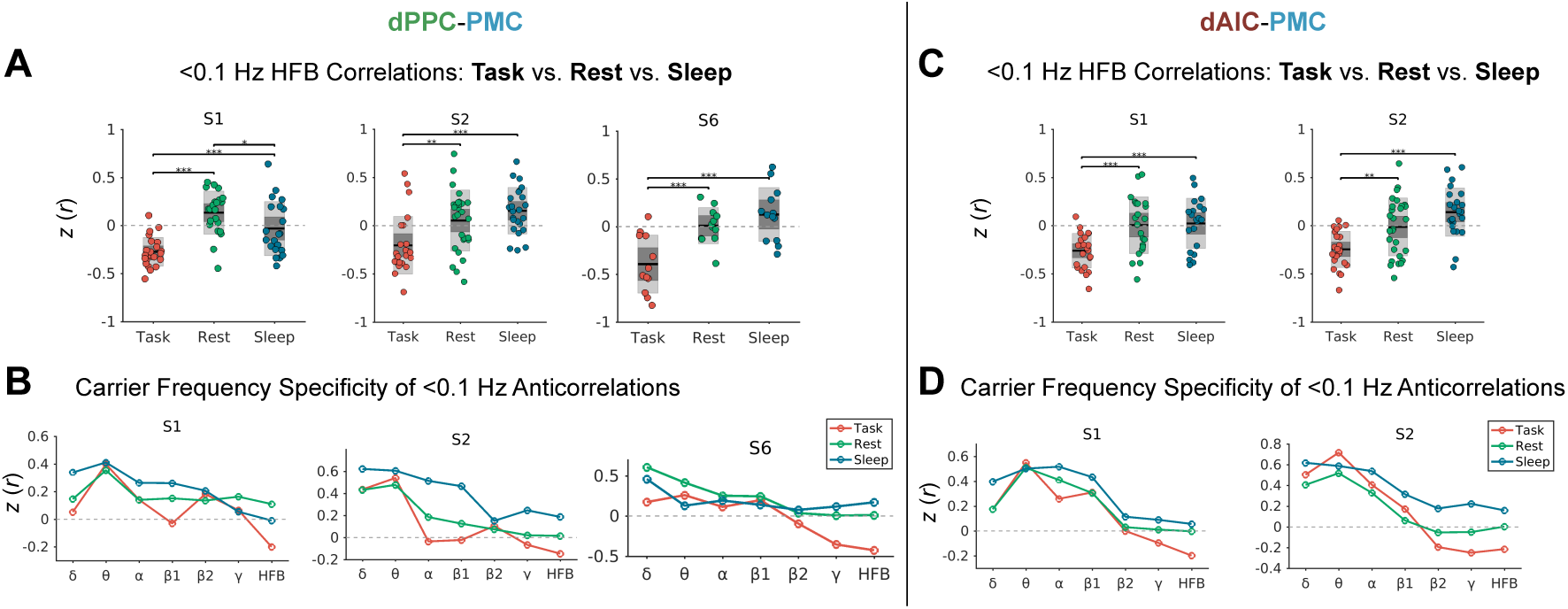
Infraslow HFB anticorrelated iEEG activity varies across task, rest and sleep states. **A)** Comparison of dPPC-PMC infraslow HFB correlations in independent 100-second windows across task, rest and sleep states in 3 patients included in cross-state analyses. **B)** In 3 patients, the average dPPC-PMC infraslow correlation across task, rest, and sleep sessions is plotted as a function of carrier frequencies including δ (1-3 Hz), θ (4-7 Hz), α (8-12 Hz), β1 (13-29 Hz), β2 (30-39 Hz), γ (40-70 Hz) and HFB (70-170 Hz). **C)** Same as A) but for 2 patients with simultaneous dAIC-PMC recordings. **D)** Same as B) but for 2 patients included in dAIC-PMC cross-state analyses. *p<0.05; **p<0.01; ***p<0.001.

To assess whether infraslow anticorrelated activity varied as a function of carrier frequency, we compared HFB with lower frequency ranges within subjects. We found that task anticorrelation was strongest and most consistent in the HFB range for both dPPC-PMC (**Figure 3B**) and dAIC-PMC (**Figure 3D**) channel pairs. Lower frequency ranges instead showed positive, rather than negative, correlations between regions across task, rest and sleep states.

### Task-like topographic network patterns during spontaneous iEEG anticorrelations

It is well-established that the spatial topography of intrinsic functional connectivity shows a high degree of stability across task, rest and sleep states in fMRI (Cole et al., 2014, Gratton et al., 2018) and iEEG (He et al., 2008, Ramot et al., 2013, Foster et al., 2015, Kucyi et al., 2018a). However, network-level topographic patterns also show spontaneous changes across various time scales, suggesting that the brain continually shifts across various ‘states,’ and some of the states could show more task-like activity patterns than others (Deco et al., 2013). As we found that the presence and magnitude of anticorrelated activity was variable across time windows in rest and sleep, we hypothesized that network-level topographic patterns would be most similar to task-like patterns when spontaneous anticorrelated activity was increased. This could suggest that spontaneous anticorrelations signify the emergence of vigilant ‘task-like’ states.

We first confirmed that a relatively stable spatial topography of activity interactions was present across all states. To do so, we performed a wider network-level analysis of correlated infraslow HFB activity from functionally localized dPPC, PMC and dAIC channels (seed regions) to *all* other implanted electrode contacts *(target* regions) within a given subject’s brain (excluding channels that were immediately neighboring the seed, were deemed pathological, were in white matter, or contained artifacts; see **Methods**) (**Figure 4A**). We then performed spatial correlations of infraslow HFB functional connectivity among iEEG task, rest and sleep states (averaged across sessions within each state) and between iEEG (all states) and within-individual resting-state fMRI connectivity patterns.

**Figure 4.**
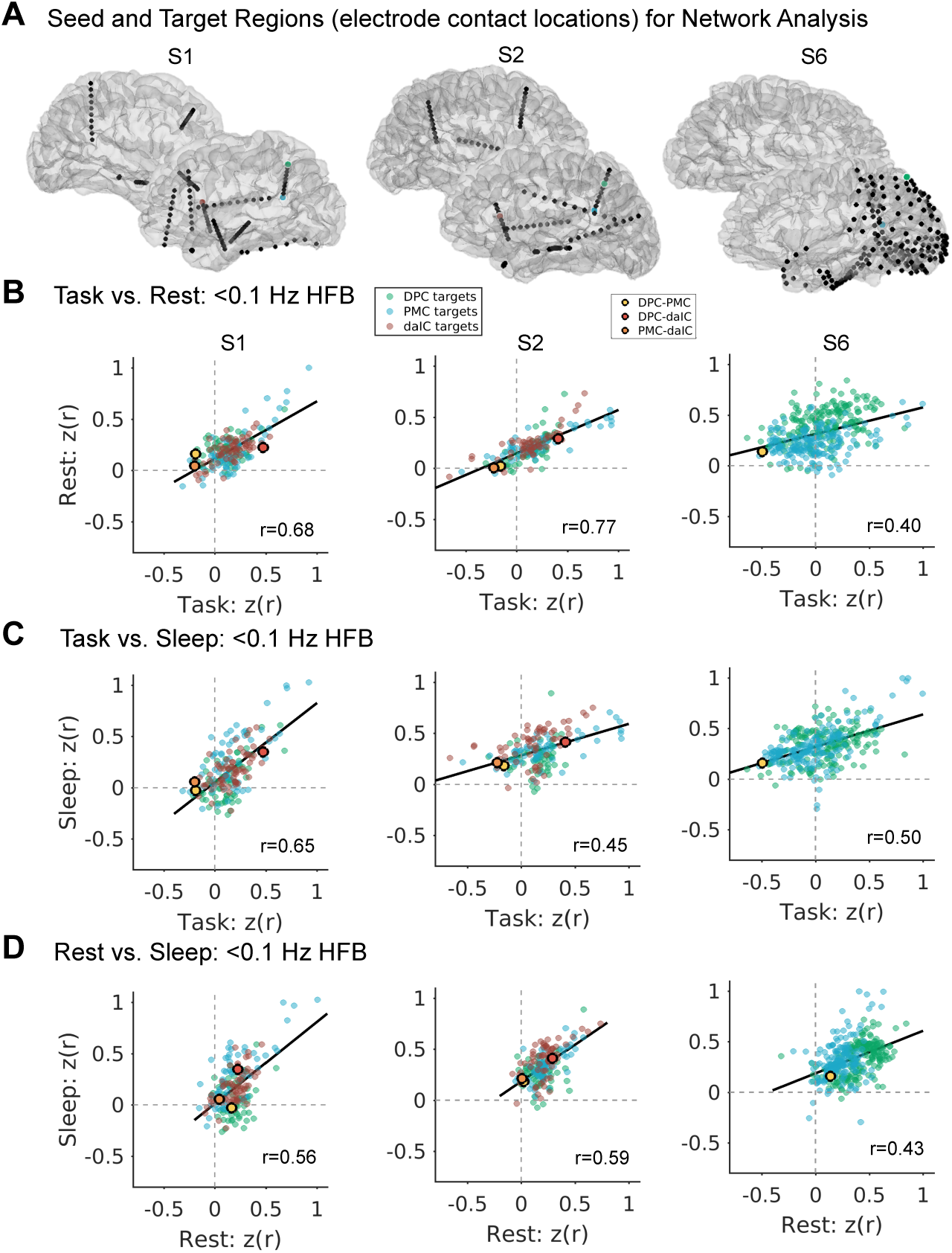
Similar iEEG infraslow HFB network topography across task, rest and sleep states. A) Locations of seed regions (green for dorsal posterior parietal cortex, blue for posteromedial cortex, red for dorsal anterior insular cortex) and other implanted electrode locations (black) in three subjects included in cross-state wider network-level analysis. B) Spatial correlations in 3 subjects of infraslow HFB functional connectivity values (Fisher-transformed) for all pairs of regions in task versus rest. C) Same as B) but for task versus sleep. D) Same as B) but for rest versus sleep. Green, blue and red data points, respectively, indicate paired targets with the dPPC, PMC and dAIC. Black-outlined yellow, red and orange data points indicate values for dPPC-PMC, dPPC-dAIC, and dAIC-PMC pairs.

This analysis revealed that the overall spatial topography of functional connectivity was similar across states (Task vs. Rest *r*=0.62 ± 0.21, all *p*<0.001; Task vs. Sleep *r*=0.58±0.09, all *p*<0.001; Rest vs. Sleep *r*= 0.58±0.09, all *p*<0.001) (**Figure 4B, 4C 4,D)**. In task, rest and sleep states, dPPC-PMC and dAIC-PMC functional connectivity values were consistently within the bottom 50 percentile (and typically within bottom 20 percentile) when compared with all other region pairs, whereas dPPC-dAIC values were within the top 50 percentile (and typically within top 80 percentile; see highlighted data points in **Figure 4**). In addition, within-individual pair-wise infraslow resting-state BOLD functional connectivity was similar to infraslow HFB iEEG functional connectivity measured in all three behavioral states (BOLD vs iEEG Task: *r*=0.41 ± 0.13, all *p*<0.001; BOLD vs. iEEG Rest: *r*=0.34 ±0.12, all *p*<0.002; BOLD vs. iEEG Sleep: *r*=0.36 ± 0.15, all *p*<0.01) (**Figure S5**). Thus, our results are consistent with the notion that intrinsic functional connectivity remains relatively stable across states despite the presence of cross-state changes in the magnitude of inter-network anticorrelation.

We next tested whether spontaneous temporal fluctuations in the degree of inter-network anticorrelation was associated with variation in the ‘task-like’ quality of topographic network patterns. In each 100-second window in rest and sleep, we determined the magnitude of dPPC-PMC or dAIC-PMC infraslow HFB anticorrelation, and we extracted the *all-to-all* connectivity matrix for implanted electrodes. We then compared the similarity of each window’s matrix with a ‘task template’ matrix that was constructed based on the average all-to-all connectivity pattern during continuous task performance (**Figure 5A**). This analysis revealed that temporal windows at rest with greater rest-task topographic similarity were associated with greater resting state anticorrelation between dPPC and PMC (β=−0.45, *t*=−4.04, *p*=0.0001) as well as between dAIC and PMC (β=−0.39, *t*=−2.97, *p*=0.005) (**Figure 5B**). Similarly, temporal windows during sleep with greater sleep-task topographic similarity were associated with greater sleep anticorrelation between dPPC and PMC (β=−0.23, *t*=−1.79, *p*=0.08) as well as between dAIC and PMC (β=−0.50, *t*=−3.81, *p*=0.004). These findings suggest that externally-oriented task-like network states emerge spontaneously during rest and sleep when inter-network anticorrelation is increased.

**Figure 5.**
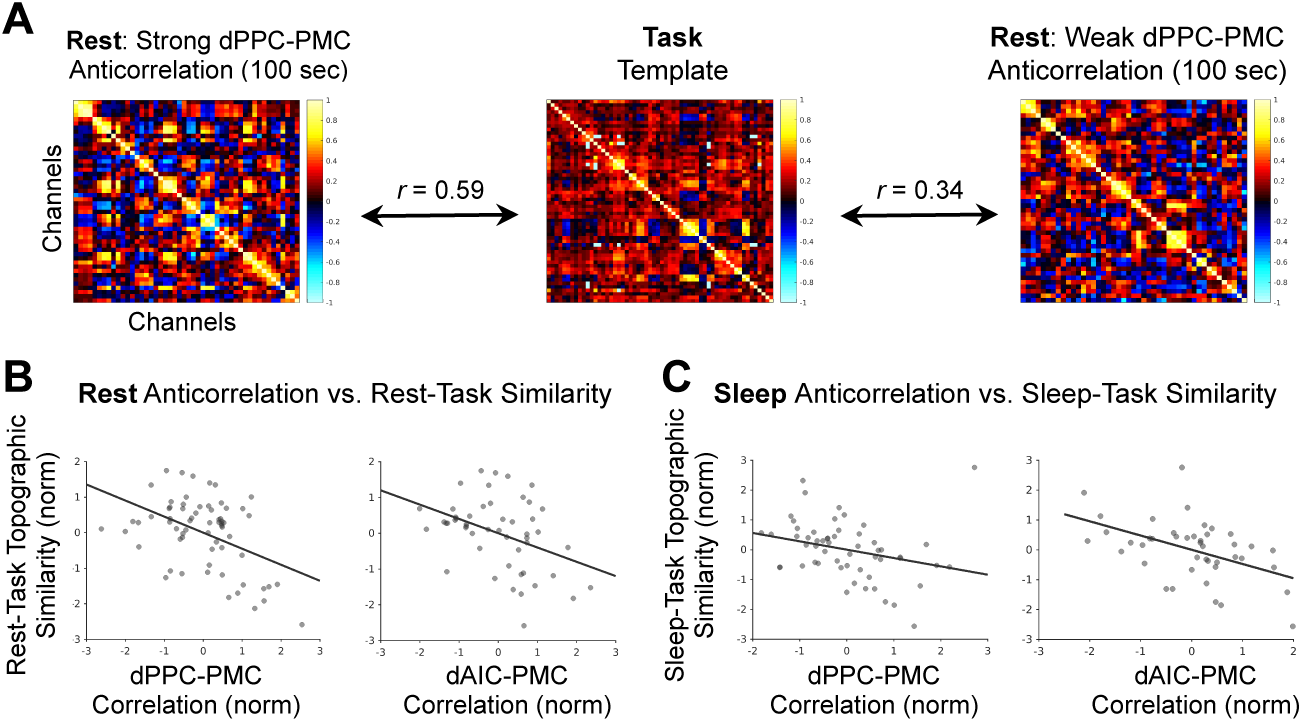
Topographic rest-task and sleep-task similarity covaries with the strength of inter-network anticorrelation. A) Example (from subject S2) channel by channel matrices of <0.1 Hz HFB correlations in 100-second resting state temporal windows with strong (left) and weak (right) dPPC-PMC anticorrelation. Compared to the window with weaker anticorrelation, the window with stronger anticorrelation shows relatively greater topographic similarity with the task template (i.e., the average correlation matrix derived from continuous task performance). B) Across independent 100-second resting state windows, a negative correlation is found between rest inter-network (dPPC-PMC and dAIC-PMC) <0.1 Hz HFB anticorrelation and rest-task topographic similarity (values normalized within subjects and concatenated across subjects). C) Same as B) but for sleep instead of rest.

### DAN and SN activations precede DMN deactivations during task performance

Our findings so far, inspired by previous fMRI studies, have considered zero-lag interactions of slow (filtered) activity fluctuations between regions. However, on a finer time scale that is measurable with iEEG, it is possible that inter-network anticorrelated activity is better explained by time-lagged interactions (Raccah et al., 2018). We therefore next sought to determine whether there was a systematic temporal order of iEEG task-evoked HFB responses across networks and whether lagged interactions could be relevant to fluctuations in GradCPT behavioral performance.

Focusing first on participants with simultaneous dPPC-PMC coverage (n=6), we found that the HFB activations in dPPC were significantly earlier than HFB deactivations in the PMC (**Figure 6A**). These responses were seen at the individual level during correct omissions but were largely absent during correct commissions (**Figure 6B**). In all six participants with relevant coverage, the HFB time-to-peak (TTP) after trial onset for maximum amplitude in the dPPC (*M±SD* = 772±208 ms) was consistently earlier than that for the peak deactivation in the PMC (1106±222 ms) (**Figure 3C**), and this timing difference was statistically significant (*p*=1.5×10^−6^, Wilcoxon signed rank test on binned subsets of trials). In three of these participants with additional electrode coverage in visual cortical areas, increased HFB power was earlier than those found for the dPPC and PMC (**Figure S3B**). Thus, the cross-regional temporal profile was consistent with an expected pattern of information transfer from unimodal to transmodal cortex, with the additional novel finding of a temporal hierarchy within the transmodal cortices themselves (i.e., the DMN deactivations hallmarking the latest stage of processing).

**Figure 6.**
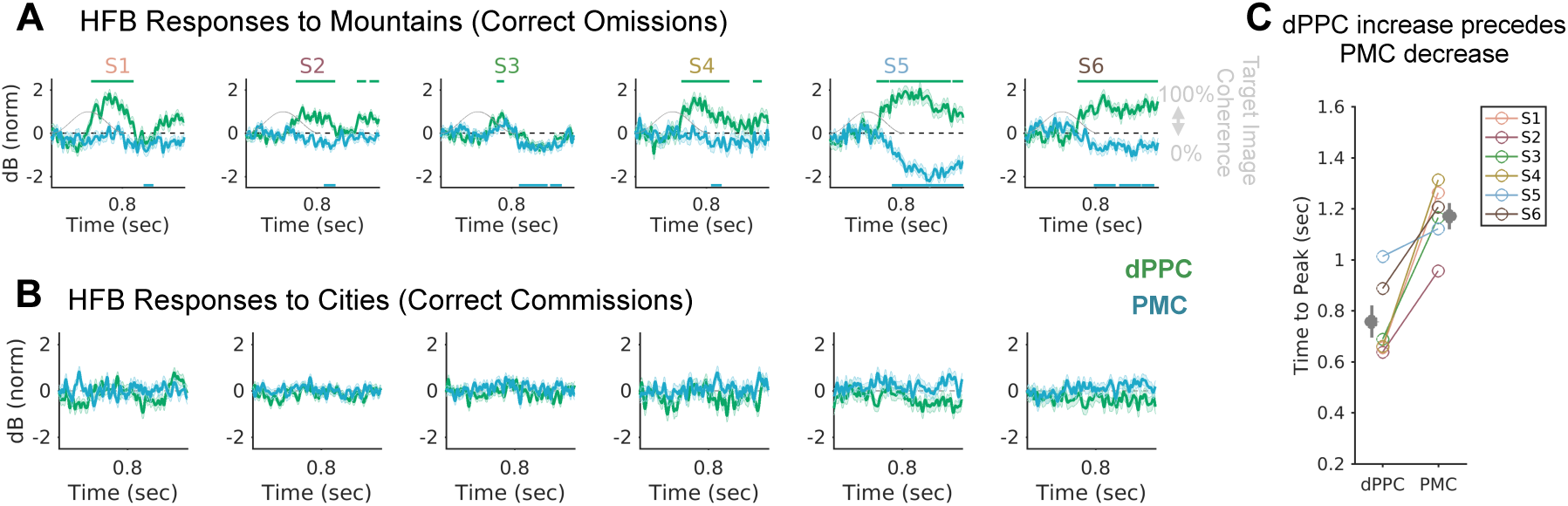
Timing of task-evoked dorsal posterior parietal cortex (dPPC) relative to posteromedial cortex (PMC) HFB responses. **A)** In all 6 participants with simultaneous dPPC-PMC coverage, dPPC peak-responsive channels showed temporal clusters of significant HFB power increases during correct omission trials (green horizontal lines above plots; Monte Carlo p<0.05), and PMC peak-responsive channels showed clusters of significant HFB power decreases (blue horizontal lines at bottom of plots; Monte Carlo p<0.05). **B)** Evoked HFB responses during correct commissions were weaker than those during correct omissions (A). C) The time-to-peak of the HFB power time course for the dPPC increase was earlier than that for the PMC decrease in all 6 subjects.

We next assessed the temporal dynamics of dAIC-PMC interactions for the 4 subjects in whom we had obtained simultaneous dAIC-PMC recordings. Similar to dPPC-PMC findings, dAIC channels illustrated earlier HFB activations (*M±SD* TTP: 813±112 ms) compared to PMC deactivations (1098±208 ms) during correct omissions (mountain) trials (p=5.1×10^−6^, Wilcoxon signed rank test) (**Figure 7A,C**). The dAIC responses were relatively attenuated during correct commission (city) trials (**Figure 7B**). Three subjects (S1, S2, S3) overlapped between the dAIC and dPPC cohorts (i.e., had simultaneous dAIC-dPPC coverage), and in these subjects, the HFB responses of dAIC compared with dPPC did not show a consistent inter-regional timing difference across subjects (**Figure S3A**).

**Figure 7.**
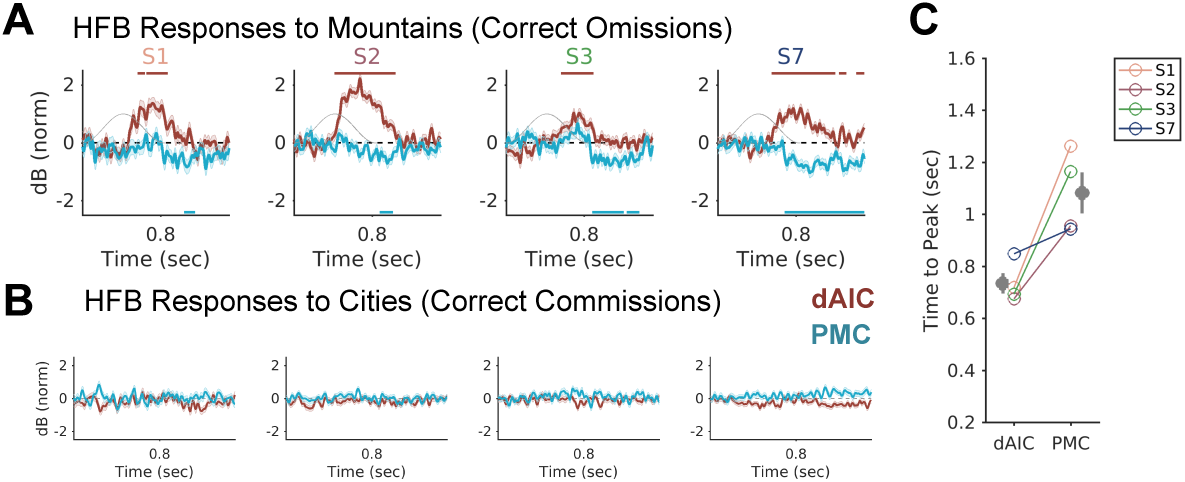
Timing of task-evoked dorsal anterior insular cortex (dAIC) relative to posteromedial cortex (PMC) HFB responses. **A)** In all 4 patients with simultaneous dAIC-PMC coverage, dAIC peak-responsive channels showed temporal clusters of significant HFB power increases during correct omission trials (red horizontal lines above plots; Monte Carlo p<0.05), and PMC peak-responsive channels showed clusters of significant HFB power decreases (blue horizontal lines at bottom of plots; Monte Carlo p<0.05). **B)** Evoked HFB responses during correct commissions were weaker than those during correct omissions. **C)** The time-to-peak of the HFB power time course for the dAIC increase was earlier than that for the PMC decrease in all 4 subjects.

Further analysis of trial-by-trial cross-correlations of inter-regional HFB time series during correct omission trials (i.e., withholding button presses to infrequent target mountain trials) revealed dPPC-PMC and dAIC-PMC anticorrelations that were *non-zero-lag* i.e., the anticorrelation between the two structures was *shifted* in time with a significant lead time from dPPC and dAIC to PMC (**Figure S4A**). A notable exception was seen in patient S3 (possible explanation provided when we discuss the behavioral significance of shifted anticorrelations). Interestingly, non-zero-lag anticorrelations were also seen, to lesser degree, for correct commission trials - i.e., frequent button presses during non-target city trials (**Figure S4A**), suggesting that temporal coordination between networks was not purely a product of responses to infrequent target stimuli.

### Behavioral significance of shifted anticorrelations between DMN and DAN

Previous fMRI evidence has suggested that time windows of greater zero-lag DMN-DAN anticorrelation are associated with better behavioral performance within and across individuals (Kelly et al., 2008, Thompson et al., 2013, Wang et al., 2016, Rothlein et al., 2018). As our time-resolved iEEG analyses had revealed a shifted (i.e., non-zero-lag) inter-network anticorrelation, we aimed to determine whether these activity lags were similarly behaviorally significant. We hypothesized that *time-shifted*, but not zero-lag, dPPC-PMC anticorrelation would reflect a subject’s level of overall sustained attention that varied across sessions.

To assess the relationship with sustained attention performance (*d*′), we estimated dPPC-PMC and dAIC-PMC functional connectivity across each task performance session using HFB envelope fluctuations. We expected that behaviorally significant time-lagged interactions would be most apparent in unfiltered, or minimally filtered, but not infraslow signals. We therefore repeated our analyses with unfiltered, 0.1-1 Hz filtered, and <0.1 Hz filtered HFB signals. The 0.1-1 Hz filter range was determined on the basis of our own and others’ prior work linking BOLD functional connectivity with HFB power amplitude in this frequency range (Nir et al., 2008, Keller et al., 2013, Foster et al., 2015, Kucyi et al., 2018a). We computed both zero-lag correlations and lag-minimum correlations, defined as the maximum anticorrelation between regions for time series that could be shifted from -2 to +2 seconds (**Figure 8A**).

**Figure 8.**
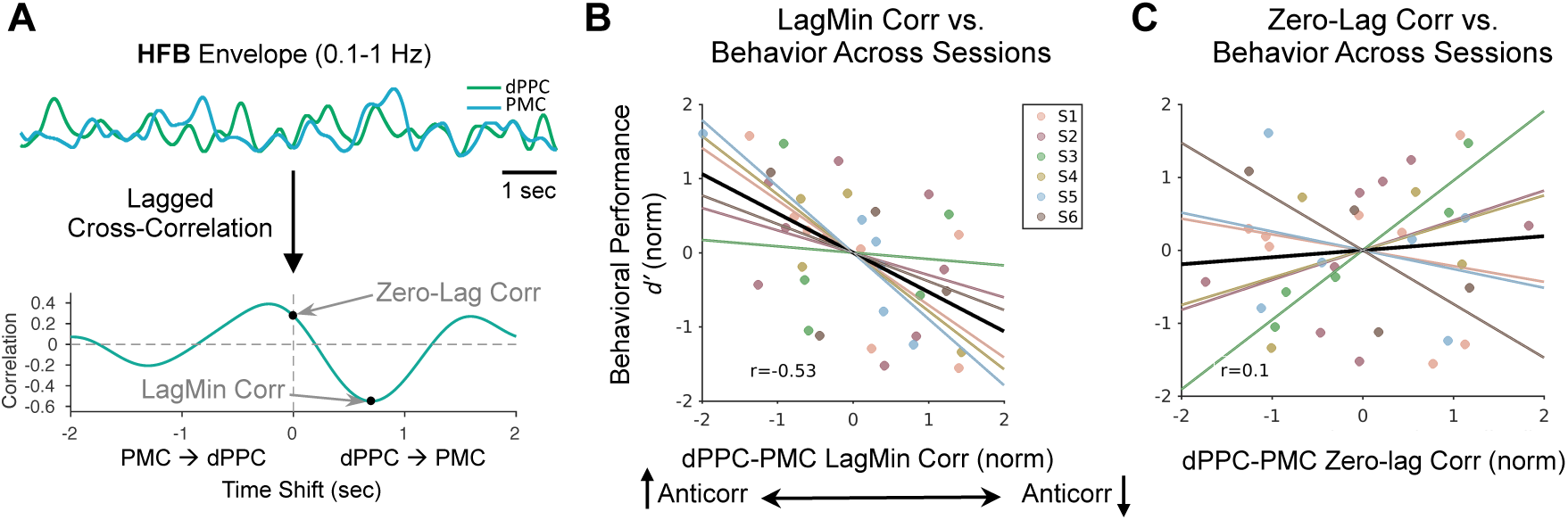
Association between time-lagged inter-network anticorrelated activity and behavioral performance across sessions. **A)** Illustration of how functional connectivity was calculated from continuous HFB 0.1–1 Hz filtered time series. Using an example 20-sec time series (top), a lagged cross-correlation was performed (shifting dPPC relative to PMC and vice versa). (Bottom) The zero-lag correlation was taken as the value with no time shift, whereas the lag-minimum correlation was taken as the minimum value across time shifts. In the main analysis, these metrics were calculated based on whole sessions (typically 6 mins long). **B)** Within-subject normalized dPPC-PMC lag-minimum correlation of 0.1–1 Hz HFB signals versus behavioral performance (d’) across sessions. **C)** Within-subject normalized dPPC-PMC zero-lag correlation of 0.1–1 Hz HFB signals versus behavioral performance (d′) across sessions. In B) and C), different colors and corresponding regression lines are plotted for individual subjects.

We found that *d*′ was significantly associated with dPPC-PMC lag-minimum correlation (β_1_=−0.56, *t*=−3.78, *p*=0.0006), but not with zero-lag correlation (β_2_=0.18, *t*=1.25, *p*=0.22), across all subjects and sessions for the 0.1–1 Hz HFB range. Specifically, greater dPPC-PMC lag-minimum correlation was associated with better sustained attention (higher *d*′) (**Figure 8B**), whereas zero-lag anticorrelation was not (**Figure 8C**). The direction of this effect was consistent within each patient, with a weakest effect for patient S3. The relationship between *d*′ and dPPC-PMC lag-minimum correlation also remained significant even when using the unfiltered, rather than 0.1-1 Hz filtered, HFB envelope (β_1_=−0.53, *t*=−2.44, *p*=0.02), suggesting consistency across multiple time scales of HFB activity. However, as expected, the association was not significant for the infraslow HFB envelope (β_1_=−0.29, *t*=−0.47, *p*=0.64; β_2_=0.32, *t*=0.52, *p*=0.61). As the distinct response profile of the dPPC in patient S3 compared to the others (**Figure 6A**) may have suggested that the recording in this subject could be from a parietal neuronal population that is functionally distinct, we repeated the analyses with S3 omitted. After omitting S3, the strength of the behavioral performance relationship with the dPPC-PMC lag-minimum correlation was similar (β_1_=−0.59, *t*=−3.71, *p*=0.0009), and the relationship remained significant when using the unfiltered HFB envelope (β_1_=−0.57, *t*=−2.64, *p*=0.01).

In contrast to the dPPC-PMC findings, sustained attention (*d*′) was not significantly associated with 0.1-1 Hz HFB envelope dAIC-PMC lag-minimum correlation (β_1_=0.98, *t*=0.45, *p*=0.66) or with zero-lag correlation (β_2_=−0.22, *t*=−1.02, *p*=0.32) across subjects and sessions. When using the unfiltered dAIC and PMC HFB envelopes, however, the relationship of sustained attention with lag-minimum correlation was significant (β_1_=−0.46, *t*=−2.31, *p*=0.03) while it was not significant for zero-lag correlation (β_2_=0.29, *t*=1.44, *p*=0.17). As expected, there were no significant relationships between infraslow HFB dAIC-PMC interactions and behavior (β_1_=−0.29, *t*=−0.49, *p*=0.63; β_2_=0.04, *t=* 0.07, *p*=0.94). Thus, dPPC-PMC lagged anticorrelation was consistently associated with attentional performance across time scales of activity, whereas dAIC-PMC lagged anticorrelation was less consistently associated with performance.

## Discussion

Here we functionally localized key nodes of the DMN, DAN and SN within individuals via analyses of neuronal population activity and resting-state fMRI network connectivity. Focusing on iEEG recordings from key DMN, DAN and SN nodes, time-resolved activity revealed dissociable task-evoked activity profiles within each network. Infraslow dPPC-PMC and dAIC-PMC anticorrelation of HFB activity was reliably detected during continuous task performance but was diminished during wakeful rest and sleep. However, during temporal windows of spontaneous infraslow anticorrelation, a ‘task-like’ wider network-level topographic organization emerged. On a finer timescale, inter-network relationships revealed a systematic temporal order of inter-network interactions, with DAN (dPPC) and SN (dAIC) activations preceding DMN (PMC) deactivations. Supporting the functional importance of this temporal orchestration, we found that greater lagged, but not zero-lag, anticorrelation between dPPC and PMC activity was associated with behavioral markers of better sustained attention across repeated sessions of continuous task performance. These findings suggest that attentional state dynamics are reflected in temporally-lagged inter-network (especially DAN-DMN) anticorrelated activity. Our results raise important issues regarding interpretation of the concept of intrinsic inter-network antagonism during active-task and task-free states.

### Attentional states and time-resolved anticorrelated inter-network activity

Anticorrelated task-evoked DMN-DAN/SN activity has been found in functional neuroimaging studies across an extensive variety of task conditions involving both stimulus-driven and goal-oriented attention (Shulman et al., 1997, Raichle, 2015). States of lapsing attention and mind-wandering, which are largely incompatible with sustained external task-oriented attention, have been associated with increased DMN activation, or a lack of DMN suppression (Weissman et al., 2006, Christoff et al., 2009, Kucyi et al., 2016). It has thus been proposed that anticorrelated networks may continuously compete with one another for control of shared computational resources (Sonuga-Barke and Castellanos, 2007, Anticevic et al., 2012).

Building on this framework, our finding that task-evoked DAN and SN activations precede DMN deactivations may point toward a causal chain of events that is required for successful deployment of stimulus-driven and goal-oriented attention. During a baseline (or low cognitive demand) state, the DMN maintains control over computational resources (e.g. for imagery associated with internally-oriented cognition). When behaviorally-relevant sensory information is successfully transferred to DAN/SN regions, those regions gain control over resources that the DMN previously had access to, and subsequently the DMN is actively suppressed. An important caveat is that our finding of a systematic temporal order of neuronal population responses does not necessarily imply a causal interaction between DAN/SN activation and DMN suppression, and further study on directional relationships and including other network nodes will be needed. Our results, however, do suggest that the relatively late suppression of the DMN relative to activation in other association networks suggests a temporal hierarchy that may accord well with findings that situate DMN regions as those with longest connectivity paths and furthest geodesic distance from primary sensory regions (Margulies et al., 2016).

The dynamics of anticorrelated activity in relation to task performance have previously been studied largely with group-level fMRI, focused on zero-lag interactions of slow hemodynamic signals. States of greater DMN-DAN/SN anticorrelation have been associated with greater vigilance and behavioral stability (Thompson et al., 2013, Wang et al., 2016, Kucyi et al., 2017, Rothlein et al., 2018). Across individuals, greater baseline BOLD anticorrelation has been associated with lesser response time variability (Kelly et al., 2008), fluid intelligence (Cole et al., 2012), and greater working memory capacity (Keller et al., 2015)– all of which are behavioral measures that may rely on sustained attention. Moreover, attenuated anticorrelation has been found in clinical conditions involving attentional dysfunction (Castellanos et al., 2008, Sripada et al., 2014) as well as in cognitive decline with aging (Keller et al., 2015, Spreng et al., 2016).

Our findings extend those results in multiple ways: First, we provide critical neurophysiological validation for slow DMN-DAN/SN anticorrelated activity during task-related sustained attention at the level of functionally localized *neuronal populations* and reliably so at the *individual* level. Second, we show that electrophysiological anticorrelated activity is associated with momentary changes in sustained attention within individuals. Third, and most importantly, we show that behaviorally-relevant anticorrelated activity involves inter-network lags of up to hundreds of milliseconds that are too short for fMRI to detect. Interestingly, the lagged inter-network anticorrelation magnitude was largely independent of zero-lag infraslow anticorrelation, a finding that accords well with the notion that infraslow brain activity has unique spatiotemporal dynamics compared to faster activity (Mitra et al., 2018). Given that such faster and slower time scales of activity were dissociable, our iEEG analyses may have been sensitive to behaviorally-relevant, time-resolved interactions that would not be detectable with current human neuroimaging methods. Despite the low temporal resolution of functional neuroimaging, inter-network directional and lagged interactions have long been of interest in task and resting states (Nyberg et al., 1996). Due to regional heterogeneity in blood flow dynamics (David et al., 2008), it remains an open question whether advances with accelerated neuroimaging will allow detection of temporally-ordered, anticorrelated DMN-DAN/SN activity and its variation over time.

### Electrophysiological interactions between DAN, DMN and SN

In the functional neuroimaging literature, the antagonistic relationship between the DMN and DAN – potentially highlighting a competition between internally-oriented and externally-oriented attention – has received intense focus and scrutiny. Though typically lesser emphasized, the SN, also shows negatively correlated BOLD activity with the DMN (as also found within individual subjects here) (Fox et al., 2005, Uddin et al., 2009, Kucyi et al., 2012). Functional neuroimaging evidence indicates that the SN and DAN have dissociable roles in externally-oriented attention. The DAN shows domain-general activation (in tandem with DMN deactivation) during various conditions involving goal-oriented attention (Corbetta and Shulman, 2002). The SN shows activation during detection of salient external stimuli (Downar et al., 2000) and during behavioral errors in fMRI (Neta et al., 2015) and single-unit recordings from key SN nodes (Fu et al., 2018). It has been proposed that the SN, and the dAIC in particular, causally facilitates switching between other networks (including DMN and DAN) to reorient attention during salient event detection (Menon and Uddin, 2010, Uddin, 2015). In partial agreement, a recent application of dynamic causal modeling to resting state fMRI data suggested that the SN and DAN exert intrinsic inhibitory influences on the DMN (Zhou et al., 2018).

Our iEEG results extend these frameworks and confirm the presence of electrophysiological DMN-SN and DMN-DAN anticorrelation during continuous task performance. The timing of task-evoked activity between DAN and SN regions was not clearly distinguishable, but activation of regions within both networks preceded DMN deactivation. Thus, our results are compatible with the possibility that the DAN and/or SN could have causal influences on DMN suppression.

We found strong evidence for dissociable electrophysiological activity in the DAN and SN. First, compared to the dPPC (DAN), the dAIC (SN) was more likely to show increased activation during behavioral errors, in line with previous fMRI evidence. Second, DMN-DAN, compared with DMN-SN lagged anticorrelation, was more strongly associated with attentional performance. Interestingly, fMRI evidence indicates that the SN may flexibly couple with either the DAN or DMN based on task conditions (Sestieri et al., 2014). Though our results here are based on an externally-oriented continuous performance task, future iEEG studies exploring distinct cognitive processes may provide further insight into context-dependent temporal dynamics of DAN, DMN, and SN interactions.

### Intrinsic inter-network anticorrelation

The fMRI-based finding of anticorrelated networks during wakeful rest has led to the notion that there is an intrinsic, state-independent, antagonistic relationship between the DMN and other networks (Fox et al., 2005). Under this framework, the brain may continuously shift between states that draw, respectively, from internally- and externally-oriented sources of information (Buckner et al., 2008, Honey et al., 2018). However, the concept has remained controversial, in large part due to technical limitations of fMRI (Murphy and Fox, 2017). Infraslow resting state BOLD anticorrelations become introduced into data following preprocessing with global signal regression (Murphy et al., 2009), but anticorrelations have also been detected in the absence of global signal regression and with alternative noise-correction strategies (also found within our cohort) (Fox et al., 2009, Chai et al., 2012). Though anticorrelation of infraslow inter-network activity has been recovered in computational models (Deco et al., 2009), electrophysiological DMN-DAN/SN anticorrelations are not often reliably observed in non-invasive M/EEG (de Pasquale et al., 2010, Brookes et al., 2011, Hipp et al., 2012). However, using 3-6 minute resting state iEEG recordings, Keller et al. (2013) showed that a subset of region pairs with resting BOLD anticorrelations exhibited anticorrelated 0.1-1 Hz HFB activity (of smaller magnitude compared to those found in BOLD data).

We investigated the electrophysiology of inter-network anticorrelation with several advances: 1) We studied functionally localized DMN-DAN and DMN-SN region pairs that were identified based on task-evoked responses; 2) We studied extended (i.e., 24-48 minutes) recordings across task, rest and sleep states; and 3) We investigated infraslow HFB activity, which closely resembles the typical temporal scale of BOLD activity studied during wakeful rest. Based on these approaches, we reliably observed anticorrelated activity during continuous task performance but not during wakeful rest and sleep states. These findings are in line with fMRI findings showing that DMN-DAN anticorrelation depends on cognitive state (Dixon et al., 2017) and is reduced during sleep (Horovitz et al., 2009, Larson-Prior et al., 2009). However, the absence of resting state iEEG anticorrelation in most of the temporal windows investigated raise questions about the validity of the notion of intrinsic inter-network antagonism.

Importantly, our findings do not conclusively rule out the possibility that intrinsic antagonistic relationships exist in the brain. We focused here on network nodes where electrode coverage was available, often only within a few sites per subject (especially in depth electrode cases). Functional heterogeneity is found across different sites within the DMN (Andrews-Hanna et al., 2010, Daitch and Parvizi, 2018), and inter-network anticorrelation may be region-dependent (Chen et al., 2017, Dixon et al., 2017).

Consistent with previous studies (He et al., 2008, Keller et al., 2013, Ramot et al., 2013, Foster et al., 2015, Hacker et al., 2017, Kucyi et al., 2018a), we found that the spatial topography of functional connectivity patterns remained similar across task, rest and sleep states and between iEEG and within-individual resting state fMRI. We also found that even when dPPC-PMC and dAIC-PMC anticorrelations were not found, inter-network correlations between these pairs were typically within the lowest percentiles compared with all other region pairs. A possible explanation is that DMN-DAN/SN pairs have an increased propensity to exhibit snippets of spontaneous anticorrelated activity. Indeed, we found that rest and sleep anticorrelations were associated with the expression of ‘task-like’ network states. It is thus possible that variations in inter-network activity coordination could be dependent on cognitive state. Though some form of behavioral validation would be needed to confirm, our task findings of increased anticorrelation with greater sustained attention support this possibility. It should be acknowledged, however, that variations in functional connectivity can arise due to sampling variability (Laumann et al., 2017). There remains an ongoing debate over whether spontaneous temporal fluctuations in functional connectivity within and between networks are behaviorally significant (Liegeois et al., 2017, Kucyi et al., 2018b).

## Conclusion

Our findings establish a behavioral significance of systematic temporal lags underlying anticorrelated inter-network activity. Additionally, our work suggests that if intrinsic antagonistic inter-network relationships exist, their expression may vary across time and as a function of behavioral state. This knowledge is critical for the interpretation of task and resting state functional neuroimaging studies and for understanding the basis of changes in inter-network relationships in health, aging and disease. The temporally ordered inter-network interactions identified here point toward the possible capacity for causal influences, a topic that requires further study with neuromodulatory techniques such as direct brain stimulation.

## Acknowledgments

We thank Clara Sava-Segal and Sori Baek for assistance with data collection and Jessica Schrouff for technical assistance with analysis. This work was supported by research Grant R01NS078396 from the National Institute of Neurological Disorders and Stroke; Grant 1R01MH109954-01 from the National Institute of Mental Health (NIMH); Grant BCS1358907 from the National Science Foundation (NSF) (all to J.P.). Data collection from Beijing was supported by the National Natural Science Foundation of China (Grant No. 81771399, 81641053, 81701276). A.K. was supported by a Banting Fellowship from the Canadian Institutes of Health Research. A.D was supported by grant F32HD087028 from the National Institute of Child Health and Human Development. M.E. was supported by a Merit Review Award from the Department of Veterans Affairs Clinical Sciences Research and Development (I01CX001653).

## Author Contributions

Conceptualization, A.K. and J.P.; Methodology, A.K., A.L.D., O.R., M.E., M.Z., C.H.; Software, A.K., M.E.; Analysis, A.K.; Investigation, A.K., M.Z., J.P., B.Z., C.Z., M.Z., C.H., K.Z., J.Z. Resources, A.K., J.P., M.E., C.Z., M.Z., C.H., K.Z., J.Z. Writing – Original Draft, A.K.; Review and Editing, A.L.D, O.R., B.Z., C.Z., M.E., M.Z., C.H., K.Z., J.Z., J.P.; Visualization, A.K. Funding Acquisition, A.K., J.P., C.Z., K.Z., J.Z.

## Methods

### Subjects

Data from seven human subjects (S1-S7) who were undergoing neurosurgical treatment for refractory focal epilepsy were included in analyses reported here. Data from S1-S6 were collected at Stanford University Medical Center, whereas data from S7 were collected at Beijing Tian Tan Hospital (age range: 19–34, 4 females and 3 male, all right-handed; see **Table S1** for full demographic and other details). Subjects were implanted with intracranial electrodes (Adtech Medical Instruments) that either were depth electrodes placed stereotactically within both hemispheres (S2, S3, S7) or one hemisphere (S4), subdural electrodes arranged in grid and strip configurations over one hemisphere (S5, S6), or a mixture of both (S1, who had all depth electrodes except for one strip covering ventral temporal cortex). Electrode placement was decided based on clinical evaluation for resective surgery. Intracranial electrode monitoring took place over the course of approximately 5–10 days at Stanford and 30 days for the patient in Beijing. Subjects at all experiment sites provided verbal and written consent to participate in research. For procedures at Stanford, The Stanford Institutional Review Board approved all procedures described herein. For procedures at Beijing, the Medical Ethics Committee of Beijing Tian Tan Hospital approved all procedures.

Only patients with simultaneous electrode coverage in the PMC and dPPC, and/or in the PMC and dAIC (as defined under *Anatomical Localization of Electrode Contacts*), were included in this study (see **Table S1** for number of electrode contacts within each of these regions for each patient). The seven patients included here were selected from a cohort of 32 patients (10 at Stanford, 22 at Beijing) who participated in the cognitive task procedures described herein. Out of those patients, 10 had simultaneous coverage in the regions of interest. Out of those 10, 3 were excluded for the following reasons: 1) irregular signals in the PMC (according to criteria defined under *Intracranial EEG: Data Preprocessing*); 2) encephalomalacia found in the occipital lobe; and 3) failure to record signals from a region of interest due to hardware failure.

### Intracranial EEG Data Acquisition

Intracranial EEG (iEEG) recordings were performed at bedside of the subject’s private clinical suite. For Stanford patients (S1–6), data were recorded either via a multichannel research system (Tucker Davis Technologies, Alachua FL, USA) or a clinical monitoring system (Nihon Kohden, Tokyo, Japan). For the Beijing patient (S7), all data were recorded with a Nihon Kohden system. The sampling rate was either 1000 or 2000 Hz for all task sessions and for some wakeful rest sessions, whereas the sampling rate was either or 500 or 1000 Hz for wakeful rest and sleep sessions (see **Table S2** and **Table S3**). For Stanford patients, depth electrode contacts (Ad-Tech Medical Instrument Corporation, Oak Creek, WI, USA) were cylindrically-shaped (0.86 mm diameter, 2.29 mm height) with inter-electrode spacing of 5–10mm. For subdural electrodes, contacts were circle-shaped with diameter of 2.3 mm in the exposed area of recording and inter-electrode spacing of 5–10 mm. For the Beijing patient, depth electrode contacts (HKHS Healthcare, Beijing, China) had a contact length of 2 mm, diameter of 0.8 mm, and inter-electrode spacing of 1.5 mm. During recording, the iEEG signals were referenced to the most electrographically silent channel outside of the seizure focus. The total number of electrode sites ranged from 60 to 210 (**Table S1**).

### Continuous Performance Task Sessions

The gradual-onset continuous performance task (GradCPT) (Esterman et al., 2013) was administered in multiple (range: 4 to 8) sessions in each patient, each lasting 4–8 minutes (**Table S2**). The number of sessions obtained within each patient depended on time available for research testing in the clinical environment, which varied across patients. The task was administered at bedside via a laptop (running Windows 10 Pro and Windows 8.1, respectively, in Stanford and Beijing) with its screen positioned ~70 cm from the patients’ eyes at chest level. Stimuli were presented using Psychophysics Toolbox (Brainard, 1997) in Matlab R2016b (MathWorks, Natick MA, USA). An RTBox device (Li et al., 2010) was used to send transistor-transistor logic pulses to an empty channel on the EEG montage to mark the onset times of each stimulus.

During task performance, grayscale visual images of either city or mountain scenes appeared within round frames (with white background) and gradually transitioned from one to another for the duration of the task. Each transition lasted 800 ms. Using linear pixel-by-pixel interpolation within each trial, image coherence began to gradually increase from time zero (minimum coherence) until 400 ms (maximum coherence) before gradually decreasing back to minimum coherence (at 800 ms). Scenes were presented randomly with 10% mountain and 90% city, but the same scene could not repeat on consecutive trials. In one subject (S7), two additional runs were performed with 25% mountain and 75% city rates, and these runs were included in functional localization analyses but were excluded from analyses of task performance versus neural activity due to potential variation in task difficulty. Subjects were instructed to press the space bar on the laptop upon noticing each city appearing but to withhold response when noticing a mountain appearing. Subjects were asked to perform their best and to keep going when they noticed themselves making an error. Each task session began with a 20 second baseline period in which the patient was instructed to fixate on a blurred mask stimulus (same size as scene stimuli) and get ready to begin. Subjects performed with their dominant hand, except for S4 who performed with their non-dominant hand due to discomfort of the dominant hand.

### Rest and Sleep Sessions

Resting state and sleep sessions were obtained and analyzed within three patients (S1, S2, and S6). Resting state recording sessions were obtained either with instructions (using a research system) or without instructions (using the clinical monitoring system) (see **Table S3** for summary). In instructed rest sessions, subjects were asked to relax and not think of anything in particular while either keeping eyes closed or open, including select sessions where subjects fixated on a central fixation cross on the laptop screen. To increase the total duration of resting state recordings, we reviewed video EEG during daytime hours and clipped periods (each at least several minutes long) when the subject appeared relaxed, awake and not engaged in conversation, clinical procedures, or with television/handheld electronic devices.

To obtain sleep recordings, we reviewed video EEG of recordings during nocturnal hours and clipped periods (each at least several minutes long) when the subject appeared to be asleep. A total duration was obtained that was similar to that obtained during task and resting states (see **Table S4** for summary). Though precise sleep staging data were not available, we inspected low-frequency power at iEEG channels of interest during wakeful and sleep recordings, as done previously (Foster et al., 2015). This allowed us to identify sleep-related features such as increased delta power, attenuation of alpha/theta oscillations and presence of sleep spindle-like oscillations at ~15 Hz (**Figure S5**).

### MRI Acquisition

In a pre-operative MRI session, all subjects underwent structural MRI (T1-weighted), and five participants (S1, S2, S3, S4, S6) underwent fMRI (T2*) during wakeful rest. In addition, a computed tomography (CT) scan was obtained following electrode implantation, which was used for anatomical localization of electrode contacts. For Stanford patients, neuroimaging was performed at Stanford Hospital on a 3.0 Tesla GE 750 MR system equipped with an 8-channel receive only head coil (8HRBrain). For patient S7, neuroimaging was performed at Beijing Dongzhimen Hospital on 3.0 Tesla Siemens MAGNETOM Verio system with a 32-channel head coil.

During resting state fMRI, subjects were instructed to relax and keep still for 6 minutes. Scan parameters for fMRI at Stanford were 64×64 mm matrix, 3.125 × 3.125 × 4.0 mm voxels, 200 mm field of view, 39 slices, 2 second repetition time, 77 degree flip angle, 30 ms echo time, and 180 volumes. For T1 scans at Stanford, the parameters were 256 × 256 matrix, 160 slices, 0.94 × 0.94 × 1.00 mm voxels, 240 mm field of view, 13 degree flip angle, 9.63 ms repetition time, and 3.88 ms echo time. For the T1 scan in Beijing, an MPRAGE sequence was acquired with parameters 256 × 256 matrix, 176 slices, 1.00 × 1.00 × 1.00 mm voxels, 9 degree flip angle, 1900 ms repetition time, and 2.53 ms echo time.

### Anatomical Localization of Electrode Contacts

We used the iElvis pipeline (Groppe et al., 2017) for anatomical localization of electrode contacts. First, we processed and reconstructed the T1 scan using Freesurfer v6.0.0 (*recon-all* command) (Fischl et al., 1999). We then aligned the post-implant CT image to the pre-implant T1 scan using a rigid transformation (6 degrees-of-freedom, affine mapping), and we inspected the quality of the registration. Using BioImage Suite (Papademetris et al., 2006), we manually labeled each electrode contact location on the T1-registered CT image. For subdural electrode cases (S5, S6), we then projected the electrode coordinates to the leptomeningeal surface and applied correction for post-implant brain shift, using previously described methods (Dykstra et al., 2012). In stereotactical EEG cases, minimal post-implant brain shift is expected, and thus no further adjustment was made. The electrode coordinates obtained from these approaches were used for visualization and for fMRI analyses.

### Intracranial EEG: Data Preprocessing

Data from each iEEG recording session were preprocessed similarly for task, rest and sleep sessions using an identical pipeline to that described in our previous work (Kucyi et al., 2018a). The procedures drew from tools in the Matlab-based LBCN preprocessing pipeline (https://github.com/LBCN-Stanford/Preprocessing_pipeline), SPM12 (Kiebel and Friston, 2004), and Fieldtrip (Oostenveld et al., 2011). For task runs, the recording was first cropped to retain data only within the baseline and task performance periods. Notch filtering was performed to attenuate power-line noise (band-stop between 57–63, 117–123, and 177–183 Hz for data from Stanford, and band-stop between 47–53, 97–103, and 147–153 Hz) for data from China. We then re-referenced the signal from each channel to the common average signal across all channels, with the following channel types excluded from the common average those that: a) showed pathological activity during clinical monitoring (as noted by a neurologist); b) were manually labeled as clear outliers on power spectra plots of all channels; c) had a variance greater or lesser than five times the median variance across all channels; or d) had greater than three times the median number of spikes across all channels, with spikes defined as 100 μV changes between successive samples. We then performed time-frequency decomposition using a Morlet wavelet transform with frequencies of interest log-spaced between 1 and 170 Hz (38 total values). To normalize the distributions of power amplitude estimates, for each frequency of interest, we rescaled each time sample by the log ratio of the whole session’s power amplitude time series. This rescaling step accounted for the band-specific *1/f* decline of the power spectrum (Miller et al., 2007). Subsequently, we performed averaging of power amplitude estimates within seven frequency bands, including *δ* (1–3 Hz), *θ* (4–7 Hz), *α* (8–12 Hz), *β1* (13–29 Hz), *β2* (30–39 Hz), *γ* (40–70 Hz) and HFB (70–170 Hz). We then visually inspected the HFB time series at each channel in each session, and we excluded channels that showed irregular, spikey or pathological activity (that may have been otherwise missed in our inspection/exclusion prior to time-frequency decomposition). Additionally, we manually reviewed the anatomical locations of electrodes and removed from analysis those channels that were located outside of the brain or largely within white matter. For each participant, we computed the average number of channels retained across sessions.

### Resting fMRI Preprocessing

All fMRI volumes were manually inspected for possible artifacts and head movements. The mean relative head displacement values for subjects (S1-S5) were 0.05, 0.03, 0.05, 0.03, and 0.09 mm. We preprocessed the fMRI data using previously described procedures (Kucyi et al., 2018a), drawing from tools in FSL v5.0.9 (Jenkinson et al., 2012), Freesurfer, Matlab and Python v2.7 (https://github.com/akucyi/rsfMRI_preproc_pipeline). The first 4 acquired volumes were deleted, followed by brain extraction (FSL’s BET), motion correction (FSL’s MCFLIRT) and linear registration between fMRI and T1 anatomical images (6 degrees-of-freedom). We automatically segmented the T1 anatomical image into white matter (WM), cerebrospinal fluid (CSF), and gray matter (GM) volumes using FSL’s FAST tool, and these volumes were registered to fMRI space. Subsequently, the segments were eroded to retain the top 198 cm^3^ and top 20 cm^3^ of voxels with highest probability of being WM and CSF, respectively (Chai et al., 2012). This erosion was performed to minimize contamination of GM signal within the WM and CSF volumes. We then regressed out of each voxel the following: mean global signal, mean WM signal, mean CSF signal, and 6 motion parameters obtained with MCFLIRT. Finally, spatial smoothing (6mm full width at half maximum kernel) and temporal filtering (0.01–0.1 Hz) were performed.

Importantly the above-described pipeline involved global signal regression, a procedure that remains controversial, especially in the context of anticorrelated brain networks (Murphy et al., 2009, Murphy and Fox, 2017). We therefore also present results from an alternative preprocessing pipeline including ICA-AROMA (Pruim et al., 2015) and without global signal regression. For this pipeline, we performed deletion of the first 4 volumes, brain extraction, motion, spatial smoothing (6mm full width at half maximum kernel), and nonlinear registration across fMRI, T1, and MNI152 standard spaces. We then applied independent components analysis (ICA) with FSL’s MELODIC and automatic dimensionality estimation, and components that were classified as noise were regressed out of each voxel, as described previously (Pruim et al., 2015). Bandpass temporal filtering (0.01–0.1 Hz) was then applied.

### Anatomical Classification of Electrode Contacts

We conducted region-of-interest (ROI) based analyses of the dorsal parietal cortex (dPPC), posteromedial cortex (PMC) and dorsal anterior insula (dAIC) due to their well-described memberships within the dorsal attention, default mode and salience networks, respectively. While other nodes of these networks were of interest, we focused on these three nodes in part due to practical considerations, as electrode coverage was based solely on clinical decision-making. We classified channels as being within the dPPC, PMC or dAIC based on individual-level anatomy reviewed on 3D T1 volumes and cortical surface reconstructions (see **Table S1** for summary of number of channels per ROI identified in each patient).

As both the superior parietal lobule and intraparietal sulcus (IPS) have been linked to the dorsal attention network (Corbetta and Shulman, 2002, Fox et al., 2006), we considered these adjacent areas within the parietal lobe as a single ROI, which we term dPPC. The IPS part of the dPPC was considered as the sulcus which runs along the anterior-posterior axis within lateral parietal cortex, approximately from the post-central sulcus to the transverse occipital sulcus. The superior parietal lobule part of the dPPC [Brodmann Area (BA) 7] included the parietal cortex regions medial to the IPS, extending in the anterior-poster axis from the post-central sulcus to the parieto-occipital sulcus.

We defined the PMC in a similar fashion as in previous work (Parvizi et al., 2006, Dastjerdi et al., 2011, Foster et al., 2015). The PMC included areas posterior to the post-central sulcus within the posterior cingulate cortex (within BA 23a and 23b), retrosplenial cortex (BA 29/30), and medial parietal cortex/precuneus (BA 31 and 7m). These areas were bounded by the marginal branch of the cingulate sulcus (dorsally/anteriorly) and by the parieto-occipital sulcus (posteriorly).

The dAIC was defined based on boundaries and landmarks defined previously (Ture et al., 1999, Naidich et al., 2004) to demarcate areas that corresponded largely to the agranular anterior insular zone (Mesulam and Mufson, 1982). This included the accessory gyrus of the insula, and/or portions of the anterior, middle and posterior short gyri of the insula that were superior to the inferior-most point of the short insular sulcus. These dorsal anterior subregions of the insula have been consistently linked with the salience, or cingulo-opercular, network (Seeley et al., 2007, Kelly et al., 2012).

For a supplemental analysis, we also defined higher-order visual cortical areas, including the lingual gyrus, fusiform gyrus and inferior temporal gyrus.

### Functional Localization of Task-responsive iEEG Channels

After identifying all electrode contacts within each ROI, we defined functionally responsive channels during GradCPT performance. Specifically, we assessed evoked HFB power during correct omissions (withheld behavioral responses) to rare, target trials (mountain scenes) relative to correct commissions (behavioral responses) to frequent, city trials. We also compared HFB responses during correct omission versus commission error (incorrect behavioral response) trials. Based on replicated findings from previous fMRI studies (Esterman et al., 2013, Fortenbaugh et al., 2018), and on the known association between BOLD activity and electrophysiological HFB activity (Logothetis et al., 2001, Mukamel et al., 2005, Nir et al., 2007, Hermes et al., 2012), we expected that HFB power would show an increase at dPPC and dAIC as well as a decrease at PMC during correct omissions and commission errors.

For this analysis, we minimally smoothed the HFB power amplitude time course within each session using a 50-ms Gaussian window. We then extracted the HFB time series from windows surrounding each mountain trial, with each window starting at 800 ms prior to mountain scene onset (start of fade-in) and ending at 1600 ms after the onset. We excluded from this analysis mountain trials that were preceded by other mountain trials. We also extracted time windows with the same boundaries around correct commission responses (city trials with button presses). For these correct commission trials, we extracted only those that were both preceded and followed by other city trials (and both with correct responses) to avoid potential contamination with responses evoked by rare mountain scenes. Among these retained correct commission trials, we deleted a random subset of trials within each session such that the remaining subset included a total number of trials that matched the number of correct omission trials within the same session. All GradCPT sessions were included in this analysis except for two runs in one patient (S1), where excessively poor performance was found (>75% commission error rate), suggesting that the patient may not have been attending to target events.

To assess significance of HFB responses during correct omission compared to correct commission trials as well as correct omission compared to commission error trials, we adopted a nonparametric cluster-based permutation test as implemented in Fieldtrip (Maris and Oostenveld, 2007) conducted separately for each ROI within each subject and accounting for multiple channels within each ROI. Combining trials across sessions within each subject, we performed independent samples t-tests on normalized HFB power amplitude values to compare conditions, using data from each time point ranging from time zero to +1500 ms relative to trial onset (beginning of stimulus fading in). For correct omission compared with correct commission trials, a one-tailed threshold of p=0.05 (negative tail for PMC, positive tail for dAIC and dPPC) was applied to the obtained t-values. For correct omission compared with commission error trials, a two-tailed threshold of p=0.05 was applied to the obtained t-values. Subsequently, adjacent samples exceeding the threshold were grouped together into clusters. The sum of t-values within each cluster was calculated for cluster-level statistics, and the maximum of these values was taken as the test statistic. These procedures were then repeated using the Monte Carlo method with 1000 randomizations of trials. Channels including observed clusters with a Monte Carlo significance probability less than 0.05 (one-tailed for correct omissions versus correct commissions, two-tailed for correct omissions versus commissions errors) were considered as significant. For the correct omission versus correct commission comparison, in cases where multiple channels showed significant responses within an ROI, we identified the channel that showed the cluster with the largest effect among all significant clusters (i.e., lowest significance probability value), and the peak-responsive channel within each ROI was selected for focused analyses described below. To evaluate the specificity of effects to HFB relative to other frequency ranges, we inspected spectrograms of mean responses during correct omission trials based on rescaled power amplitude estimates in the range of 1–170 Hz (**Figure 1**) (for visual purposes, smoothing with a Gaussian kernel was applied).

### Multiple Kernel Learning Analysis of iEEG Responses

To comprehensively assess the possible contributions of different frequency bands of activity to task-evoked iEEG responses, we performed a multiple kernel learning (MKL)-based analysis. The MKL approach is a machine learning method that can be applied to classifying iEEG task conditions by including multiple frequency bands of activity as well as multiple channels in a single model (Schrouff et al., 2016). We used MKL in the PRoNTo toolbox (Schrouff et al., 2013) to classify correct omission versus correct commission trials.

For each subject, all channels that were anatomically identified as being within one of the ROIs (PMC, dPPC, dAIC) were included in the model. In a *full* model, channels from all ROIs within a given subject were included, and in additional single ROI models, channels only from given ROIs were included. For each trial, power amplitudes from each channel were extracted between time 0 to 1500 ms after trial onset for seven frequency bands (*δ*, *θ, α, β1, β2, γ*, HFB). The number of trials was matched between conditions (as described above). Model features were defined as *kernels*, or pair-wise similarity matrices across the time series of all trials, which were constructed for each channel and frequency band (i.e., the number of kernels per subjects was *m* × 7, with *m* being the number of channels). Each kernel was normalized and mean-centered to ensure that modeling was not influenced by the scale of each kernel.

We then applied MKL, using a support vector machine to define a decision boundary to discriminate between correct omission and correct commission trials. As in previous work (Schrouff et al., 2016), model parameters were optimized to determine the decision boundary for each kernel, and decision boundaries were weighted by a parameter *d_m_* to define a global decision boundary. We used a 10-fold cross-validation scheme: in each fold, training was performed 90% of trials, and testing was performed on the 10% left out trials (with a different 10% left out on each fold). Model accuracy was obtained as the average balanced accuracy (average of class accuracies) across folds. During cross-validation, the soft-margin parameter, *C*, was optimized by considering values 0.01, 0.1, 1, 10, 100 and 1000. A nested cross-validation was performed where the value of *C* leading to highest model performance in the inner cross-validation was selected, and that *C* value was used to estimate performance in the outer cross-validation. Statistical significance of model accuracy within each subject was assessed using 1000 permutations of the training labels to generate an accuracy distribution, and *p* values less than 0.05 for the true value were considered as significant. To evaluate the contributions of different frequency bands of activity to model performance, for each fold we calculated the sum of *d_m_* values across channels for each of the seven frequency bands. We then calculated the mean of those sums across the ten folds.

### Resting fMRI: Seed-based Functional Connectivity

We transformed the within-subject coordinates obtained from electrode localization to fMRI space (using the previously computed linear transform). We then extracted the BOLD time series from seed regions defined as 6-mm radius spheres at electrode locations of interest (i.e., peak-responsive channels, defined based on criteria described above). Using a general linear model (implemented in FSL) for each seed region separately, the demeaned the BOLD time series was entered as a regressor. We then projected the resulting volume-map z-scores obtained at each voxel to vertices on the cortical surface in Freesufer. To provide a comparison between individual-level networks found with this approach and the standard networks found in healthy populations, we registered the DMN, DAN and SN templates from the 7-network Yeo parcellation (Yeo et al., 2011) from standard (fsaverage6) to individual surfaces.

In the two patients who did not undergo fMRI, we registered the Yeo parcellation to individual cortical surfaces to determine whether peak-responsive channels were within the DMN, DAN or SN. As one of these two patients had depth electrodes (rather than subdural surface recordings), raw electrode coordinates were in volume space rather than in Freesurfer cortical vertices. Thus we snapped the coordinates to the nearest cortical vertex and determined its Yeo network identity.

### Intracranial EEG: Comparison of Task, Rest and Sleep States

We compared HFB functional connectivity across states (task, rest and sleep) for dPPC vs. PMC and dAIC vs. PMC, using the functionally localized peak-responsive channels for each subject within ROIs. To increase the statistical power of this analysis, we divided each task, rest and sleep recording session into 100-second, non-overlapping windows (deleting the remaining last part of each recording). The 100-second window length was selected so that a slowest frequency component of 0.01 Hz, which is commonly assumed to be relevant to infraslow functional connectivity (Leonardi and Van De Ville, 2015), was retained. Within each window, we computed functional connectivity as the Fisher-transformed inter-regional correlation of the infraslow (<0.1 Hz) HFB envelope. Using these functional connectivity values, we performed a three-way analysis of variance (ANOVA) within each subject with factors Task, Rest and Sleep (significance set at two-tailed *p*<0.05). We performed post-hoc two-sample t-tests to compare functional connectivity between pairs of states and one-sample t-tests to assess whether non-zero functional connectivity was reliably detected within each state (significance set at two-tailed *p*<0.05). We also repeated the same analyses using power amplitude correlations in six other frequency bands other than HFB (*δ*, *θ, α, β1, β2, and γ*).

We additionally performed a wider network-level functional connectivity analysis to compare the spatial topography of HFB inter-regional correlations across states. For this analysis, all channels within each subject that were retained after preprocessing were included. Using the peak-responsive channels within the dPPC, PMC and dAIC as seed regions, we computed the inter-channel time series correlations (Fisher-transformed) within each session and then averaged these values across sessions for each state (i.e., generating a vector for each seed region to all target regions). To avoid potential spurious correlations due to volume conduction, we excluded target channels that were immediately neighboring the seed channel (on the same depth probe or subdural strip). We then concatenated vectors across seed regions within subjects and deleted values from redundant pairs. We then correlated the obtained vectors from the three states (task, rest sleep) with one another as an index of the inter-state spatial similarity of HFB functional connectivity (significance set at p<0.05). We also repeated these analyses of wider network-level functional connectivity using power amplitude correlations in six other frequency bands other than HFB (*δ*, *θ, α, β1, β2, and γ*).

We performed additional analyses to test the hypothesis that task-like topographic network patterns would be found during temporal windows in which rest and sleep infraslow HFB anticorrelations emerged. For this analysis, we computed a ‘task template’ topographic pattern, defined as the mean correlation matrix (Fisher-transformed and using all channels retained for analysis) across all independent 100-second windows during task performance. For each rest and sleep 100-second window, we computed the correlation matrix (Fisher-transformed) between the same channels. We then correlated each rest and sleep matrix with the task template. For the two ROI pairs of interest (dPCC-PMC and dAIC-PMC) and the two behavioral states (rest and sleep), we then performed linear mixed model analyses, as implemented in the R environment (Baayen et al., 2008). Across all subjects and temporal windows, subject was entered as a random effect, similarity to task template was entered as dependent variable, and ROI pair correlation was entered as fixed effect. For these analyses, task template similarity and ROI pair correlation values were within-subject normalized (by mean and standard deviation). Significance was set at *p*<0.05 (Sattherthwaite’s approximation) (Luke, 2017).

### Intracranial EEG versus BOLD Functional Connectivity Analysis

For each state recorded in iEEG data (task, rest, sleep) we compared the spatial topography of HFB functional connectivity with that of BOLD functional connectivity within subjects. For this analysis, we extracted the BOLD time series from 6-mm radius spheres surrounding each channel’s coordinates in fMRI space (as described above). When this procedure resulted in overlapping voxels between regions (e.g. neighboring channels), we deleted those voxels from each region before extracting the time series. Using the locations of the peak-responsive channels within the dPPC, PMC and dAIC as seed regions, we subsequently computed the inter-regional BOLD correlation (Fisher-transformed) with all target regions (excluding those regions located at channels that were excluded from iEEG analysis). We then concatenated vectors across seed regions within subjects and deleted values from redundant pairs. We then correlated the BOLD vectors with the iEEG vectors from each state (task, rest, and sleep) with regions/channels aligned across modalities (significance set at *p*<0.05).

### Intracranial EEG: Time to Peak Estimation

We performed time-to-peak (TTP) estimation of HFB responses during correct omission trials on peak-responsive channels within the dPPC, PMC and dAIC identified within each subject. Within a time window ranging from +200 to +1500 ms after trial onset (i.e., beginning of mountain scene fade-in), we identified the maximum peak time point for dPPC and dAIC and the minimum peak time point for PMC based on the average HFB response across trials (**Figure 3C**). However, because only 6, and 4, subjects were included in these analyses for dPPC vs. PMC and dAIC vs. PMC, respectively, we performed additional data processing so that more powerful statistics could be performed. Specifically, we divided all trials into equally-sized bins of trials, selecting 8 total bins per subject (i.e. the maximum number of sessions per subject). In each bin we computed TTP as described above. We entered TTP within all bins and across all subjects into two-sided Wilcoxon signed rank tests comparing a) dPPC versus PMC; and b) dAIC versus PMC (significance set at *p<0.05*).

In a complementary approach to TTP analysis, we performed cross-correlations between channels’ time series, with shifts ranging from -2 to 2 seconds. These cross-correlations were performed on single correct omission (mountain) trials, and the mean across cross-correlation across trials was plotted (**Figure S4**). We repeated this cross-correlation analysis for all correct commission (city) trials that were preceded and followed by other city trials.

### Behavioral Analysis

We used sensitivity (*d*′) as a measure of task performance, based on signal detection theory (Macmillan and Creelman, 2004), within each session:

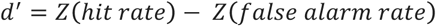

where *Z*(*p*) is the inverse of the cumulative distribution function of the Gaussian distribution. Thus, the higher the *d*′ value, the higher the overall accuracy of behavioral performance (based on responses to both cities and mountains). Evidence indicates that *d*′, based on GradCPT performance, is a generalizable measure of sustained attention (Fortenbaugh et al., 2015, Rosenberg et al., 2016).

Because of the fast pace of the task and the overlap of stimuli across adjacent trials, key presses were assigned to trials using a previously described iterative algorithm (Esterman et al., 2013, Kucyi et al., 2016). Presses were assigned relative to the beginning of each image transition. For trials in which the reaction time (RT) was highly deviant (before 70% image coherence for current trial, or after 40% coherence for the following trial), the following criteria were used for trial assignment: 1) If the previous or current trial had no response, the press was assigned to the trial in which the response occurred; 2) If both adjacent trials had no response, the press was assigned to the trial closest in time (excluding cases where the trial was a mountain image). 3) If multiple presses could be assigned to a given trial [based on 1) and 2)], the fastest RT was assigned to that trial.

### Task-based iEEG Functional Connectivity Analysis

Prior to functional connectivity analysis, we applied a bandpass temporal filter (butterworth, 4^th^ order) to the unsmoothed HFB envelope, retaining frequencies between 0.1–1 Hz (Nir et al., 2008, Keller et al., 2013, Kucyi et al., 2018a). We performed additional analyses of the HFB envelope, based on no filtering (minimally smoothed, as described above) as well as lowpass-filtering of the time series to the infraslow (<0.1 Hz) range. Task-based functional connectivity analysis was applied for peak-responsive channels in the dPPC, dAIC and PMC, using two metrics: 1) Zero-lag FC: the zero-lag correlation between the channels’ time series, and 2) Lag-minimum FC: the minimum correlation (i.e., the greatest anticorrelation) among cross-correlations between the channels’ time series, with shifts ranging from -2 to 2 seconds. We then applied a Fisher r-to-z transformation to these values.

To compare functional connectivity with session-to-session variability in behavioral performance (*d*′) across subjects, we first normalized functional connectivity and *d*′ values within subjects (i.e., for each session, we subtracted out the mean and then divided by the standard deviation of values across sessions). We then performed linear mixed effect model analyses with subjects entered as random effects, behavioral performance as dependent variable and functional connectivity (zero-lag FC and lag-minimum FC) as fixed effects. Significance was set at *p*<0.05 (Sattherthwaite’s approximation). For visual display purposes, we plotted normalized functional connectivity versus (*d*′) values, including regression lines illustrating group- and individual-level data (**Figures 3E and 3F**). Only GradCPT sessions that included 90% city and 10% mountain rate (**Table S2**) were included in these analyses so that task difficulty was matched across sessions.

## Supplemental Information

### Supplemental Figure Legends

**Figure S1.**
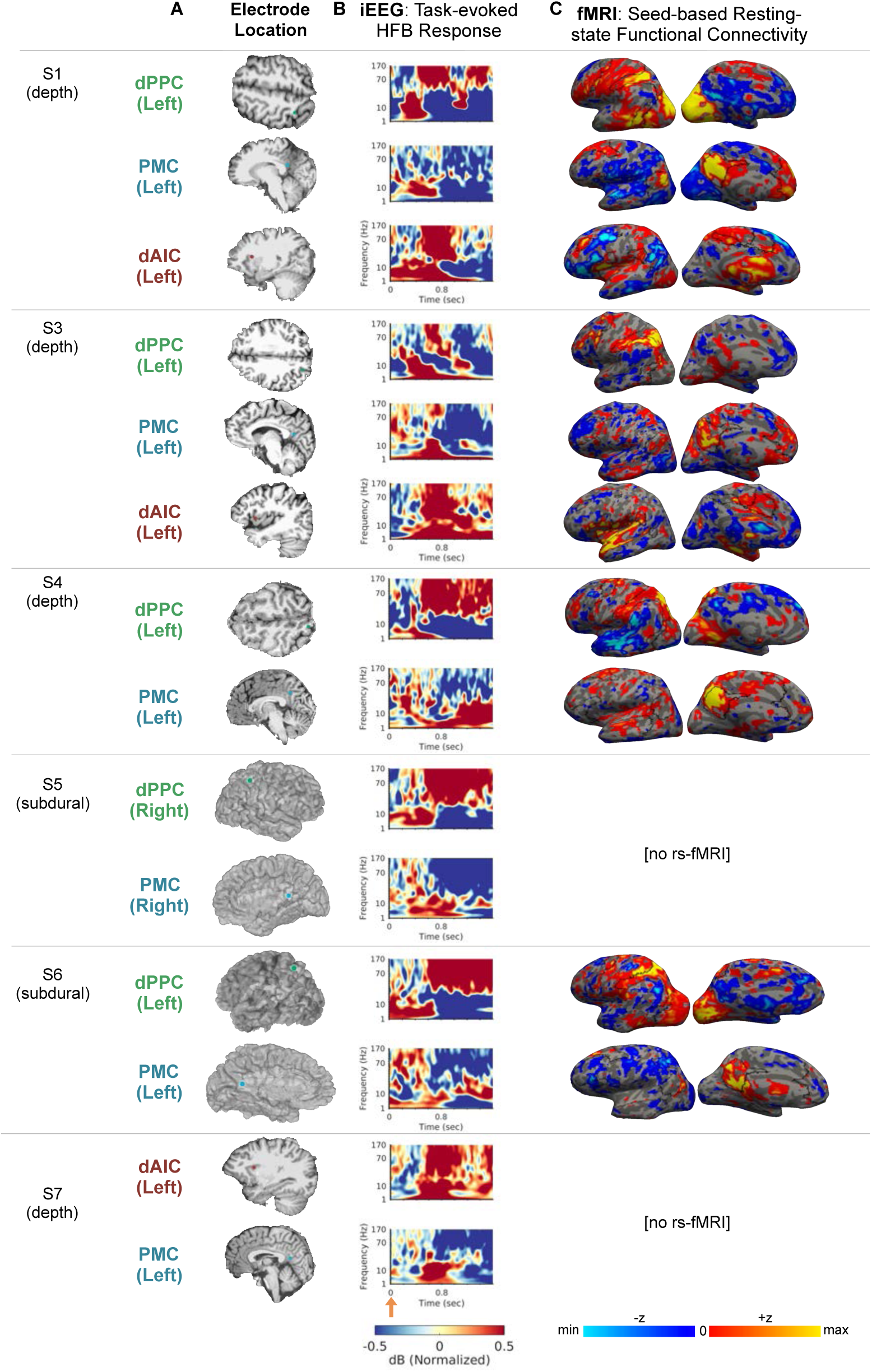
Functional localization of dPPC, PMC and dAIC in all subjects (except S2; see **Figure 1**). (Left) Anatomical locations of electrode contacts implanted in the dPPC, PMC, and dAIC that showed peak-responsive high-frequency broadband (70–170 Hz) responses during withheld responses to mountains (correct omissions). (Middle) Time-frequency plots highlighting spectral changes during correct omissions. (Bottom) Resting-state functional connectivity (based on within-individual pre-operative fMRI) from seed locations at the dPPC, PMC and dAIC electrode contacts. Red/yellow indicates positively correlated regions; blue/light blue regions indicates negatively correlated regions (z scores based a general linear model analysis thresholded arbitrarily for display purposes).

**Figure S2.**
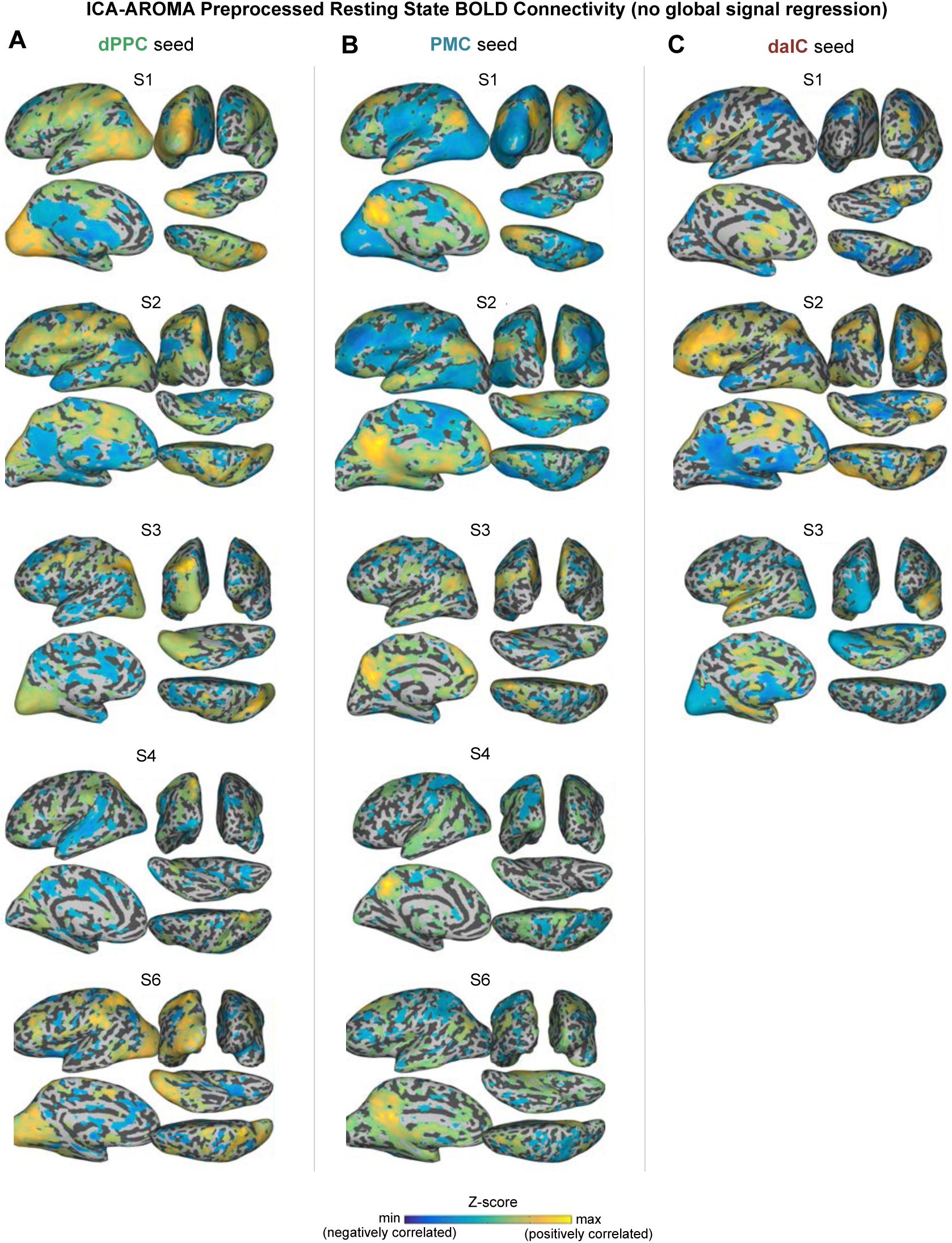
Resting-state BOLD functional connectivity (based on within-individual pre-operative fMRI) with ICA-AROMA data preprocessing. Functional connectivity maps from dPPC (left), PMC (middle) and dAIC (right) seed locations that were defined based on peak-responsive iEEG sites within individuals. Yellow indicates positively correlated regions; blue indicates negatively correlated regions (z scores based a general linear model analysis thresholded arbitrarily for display purposes).

**Figure S3.**
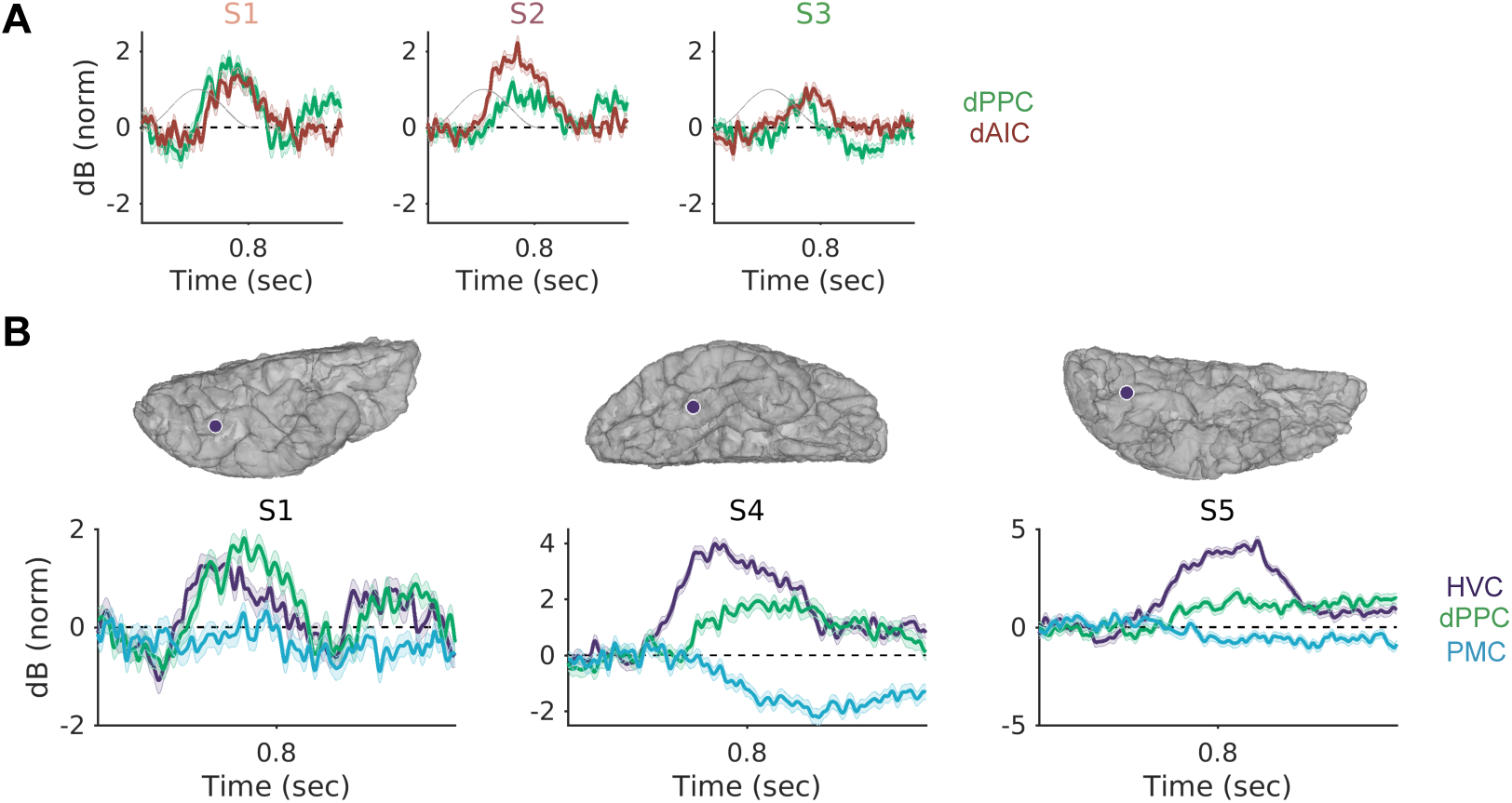
High-frequency broadband (HFB; 70–170 Hz) responses across multiple regions during correct omission trials. A) Plots showings responses of peak-responsive electrode contacts in the dPPC (green) and dAIC (red) in three subjects with simultaneous coverage in the two regions. B) Plots showings responses of areas within higher-order visual cortex (HVC; purple) relative to peak-responsive dPPC (green) and PMC (blue) sites in three subjects with simultaneous coverage in all three regions. Anatomical images (top) show locations of subdural HVC electrode contacts.

**Figure S4.**
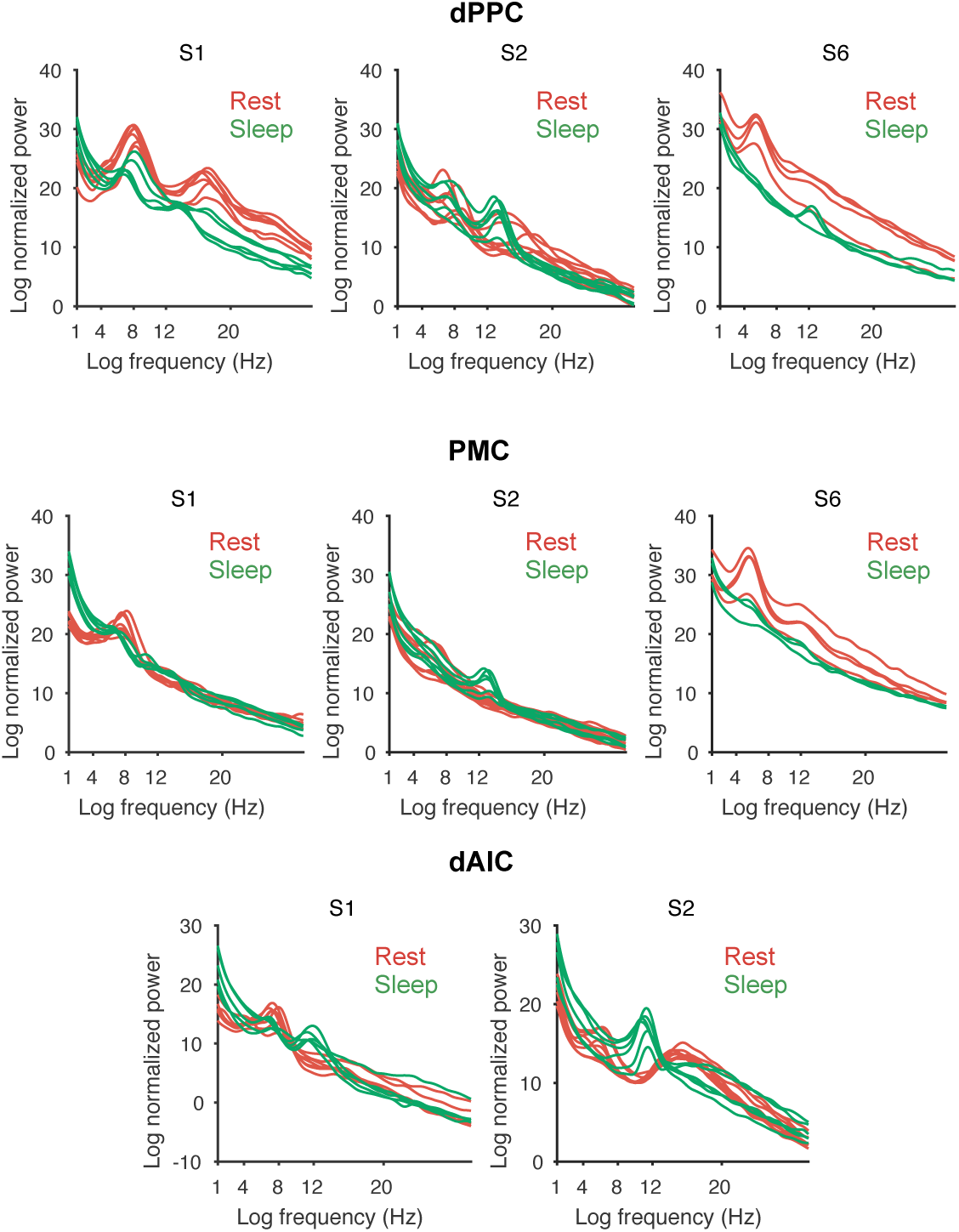
Low-frequency power spectra for wakeful rest (red) versus sleep (green) recording sessions. For each recording session (each shown as different curve), power spectra in 1–30 Hz range (plotted as log-frequency and log-normalized power) are shown for peak-responsive channels in the dPPC (top), PMC (middle) and dAIC (bottom). Power spectra were calculated from each whole-session time series using Welch’s method for spectral density estimation (fast Fourier transform length of 10 times sampling rate, 0.5 overlap between windows). In several plots, theta/alpha oscillations show attenuation in sleep compared to wakeful rest, and sleep spindle-like oscillations (~15 Hz) are seen in sleep.

**Figure S5.**
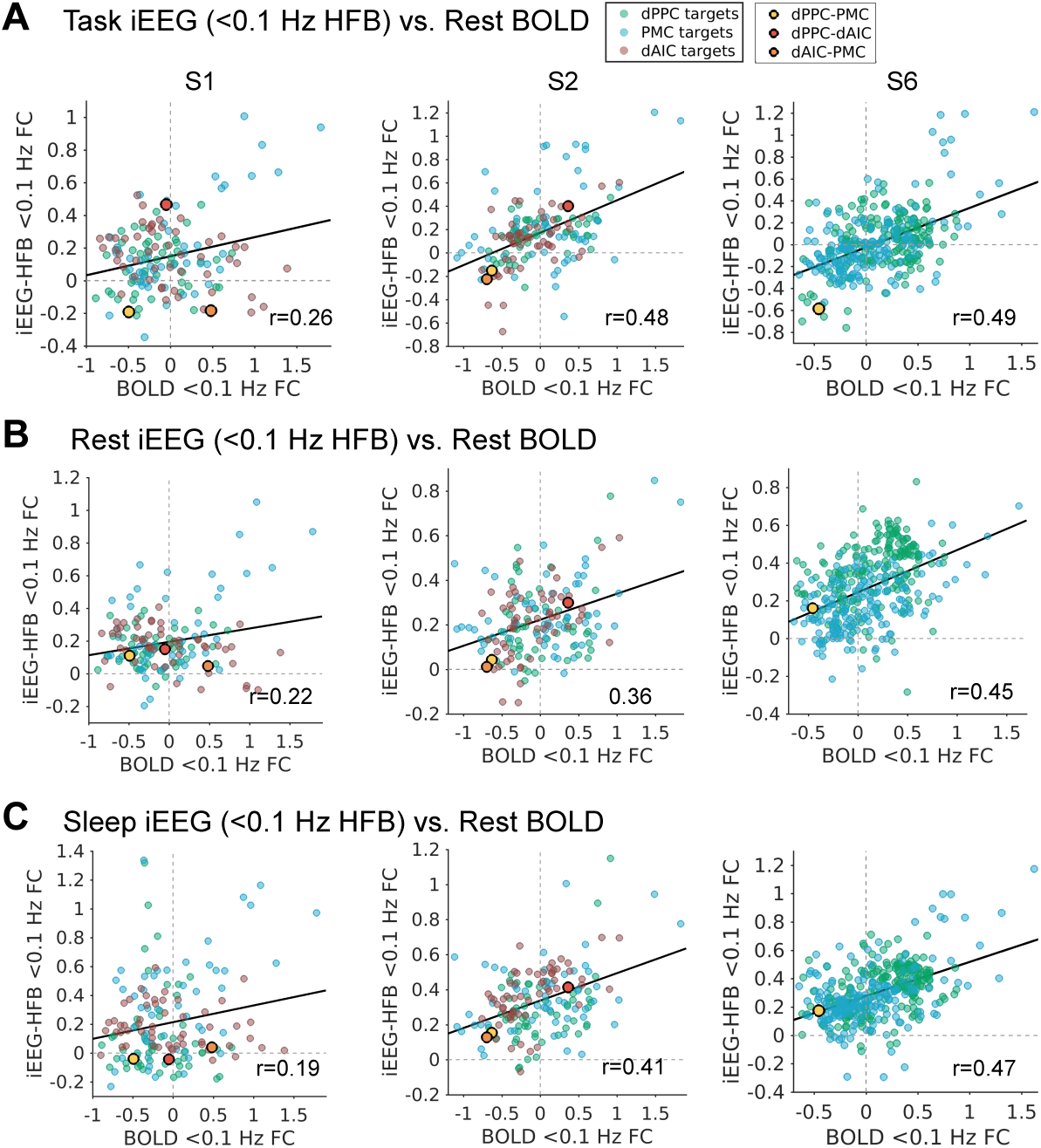
Similar iEEG compared to resting-state BOLD network topography. A) Spatial correlations in 3 subjects of task iEEG infraslow HFB versus BOLD functional connectivity values (Fisher-transformed) for all pairs of regions in iEEG task versus BOLD rest. B) Same as A) but for iEEG rest versus BOLD rest. C) Same as A) but for iEEG sleep versus BOLD rest. Green, blue and red data points, respectively, indicate paired regions with the dPPC, PMC and dAIC. Black-outlined yellow, red and orange data points indicate values for dPPC-PMC, dPPC-dAIC, and dAIC-PMC pairs.

**Figure S6.**
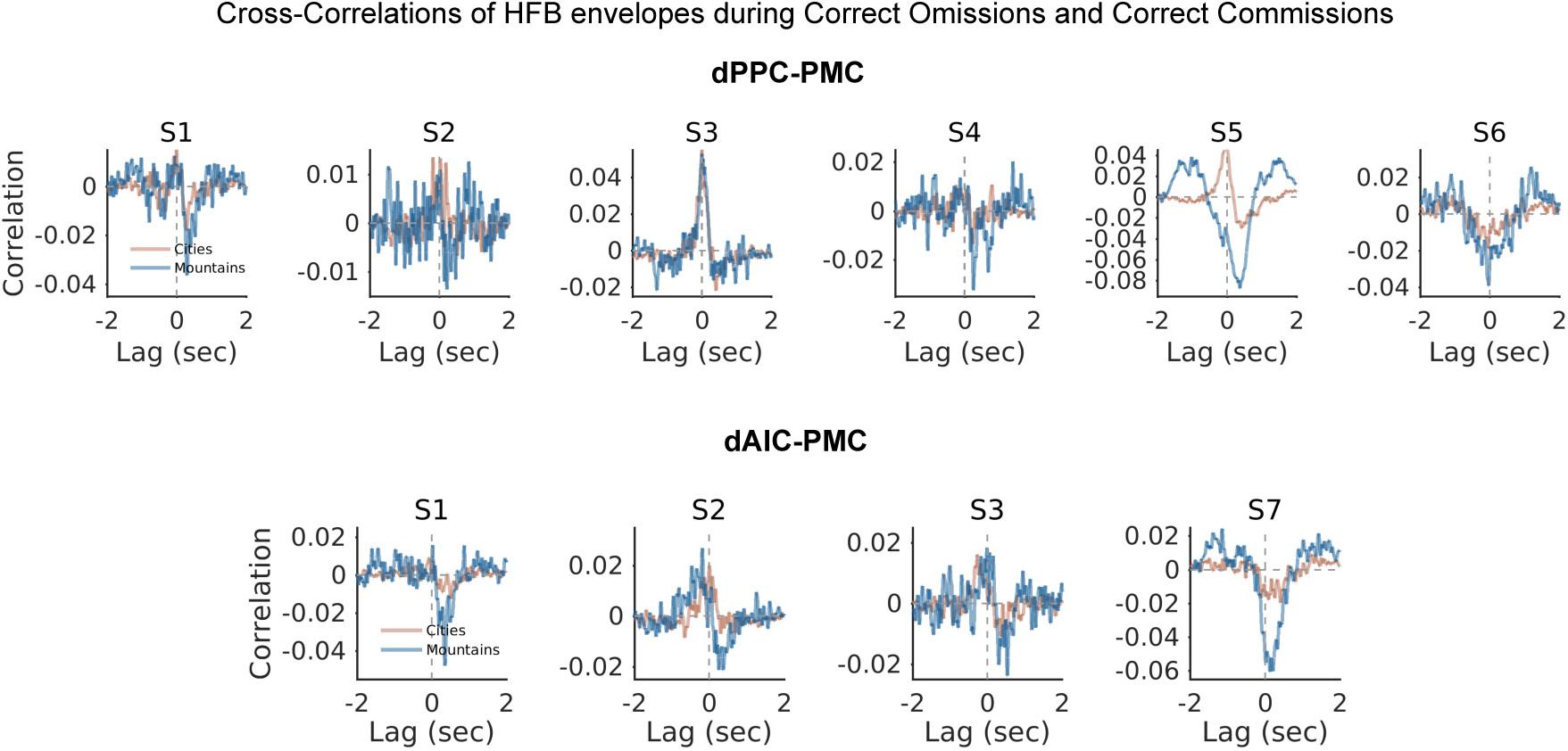
Average trial-by-trial lag cross-correlations of HFB power between dPPC-PMC and dAIC-PMC for correct omissions and correct commissions. For each trial within each category, cross-correlations were calculated between channels’ time series, with shifts ranging from -2 to 2 seconds. The mean across cross-correlation values across trials for each category is shown for 6 subjects with simultaneous dPPC-PMC coverage and 4 subjects with simultaneous dAIC-PMC coverage.

### Supplementary Tables

**Table S1.** Subject demographics and characteristics

|  | **S1** | **S2** | **S3** | **S4** | **S5** | **S6** | **S7** |
| --- | --- | --- | --- | --- | --- | --- | --- |
| **Site** | Stanford | Stanford | Stanford | Stanford | Stanford | Stanford | Beijing |
| **Subdural/Depth** | Both | Depth | Depth | Depth | Subdural | Subdural | Depth |
| **Age** | 31 | 19 | 34 | 27 | 21 | 30 | 27 |
| **Sex** | M | F | F | F | M | F | M |
| **Handedness** | Right | Right | Right | R | Right | Right | Right |
| **Implanted Hemisphere** | Both | Both | Both | Left | Right | Left | Both |
| **Epileptic Focus** | Bilateral temporal lobe | Left lateral temporal lobe | Left medial and lateral temporal lobe | Left precentral gyrus | Right posterior temporal lobe | Left occipital and angular region | Left inferior parietal lobule |
| **Duration of Epilepsy** | 7 years | 5 years | 18 years | 19 years | 8 years | 20 years | 16 years |
| **# of electrode contacts (gray matter/total)** | 76/136 | 71/116 | 62/90 | 50/60 | 128/128 | 210/210 | 94/156 |
| **# dPPC contacts (responsive/total)** | 2/2 | 3/12 | 1/5 | 2/7 | 3/4 | 4/14 | 0/0 |
| **# PMC contacts (responsive/total)** | 1/7 | 1/4 | 3/3 | 1/4 | 1/2 | 7/18 | 4/8 |
| **# dAIC contacts (responsive/total)** | 2/10 | 9/12 | 1/2 | 0/0 | 0/0 | 0/0 | 4/4 |

**Table 2.** Summary of task sessions conducted.

| Patient | Run # | Duration<br>(mins) | City/Mountain<br>Rate | $d'$ | Commission<br>Error Rate | Omission Error<br>Rate |
| --- | --- | --- | --- | --- | --- | --- |
| **S1** | 1 | 6 | 0.1/0.9 | 2.88 | 0.38 | 0.0049 |
|  | 2 | 6 | 0.1/0.9 | 3.41 | 0.35 | 0 |
|  | 3 | 6 | 0.1/0.9 | 2.86 | 0.57 | 0 |
|  | 4 | 6 | 0.1/0.9 | 2.84 | 0.49 | 0.0025 |
|  | 5 | 6 | 0.1/0.9 | 2.96 | 0.53 | 0 |
|  | 6 | 6 | 0.1/0.9 | 2.23 | 0.79 | 0 |
|  | 7 | 6 | 0.1/0.9 | 2.11 | 0.81 | 0 |
|  | 8 | 6 | 0.1/0.9 | 2.78 | 0.69 | 0 |
| **S2** | 1 | 6 | 0.1/0.9 | 2.33 | 0.28 | 0.040 |
|  | 2 | 6 | 0.1/0.9 | 2.19 | 0.27 | 0.057 |
|  | 3 | 6 | 0.1/0.9 | 2.65 | 0.32 | 0.015 |
|  | 4 | 6 | 0.1/0.9 | 2.57 | 0.34 | 0.015 |
|  | 5 | 6 | 0.1/0.9 | 3.08 | 0.40 | 0.0024 |
|  | 6 | 6 | 0.1/0.9 | 3.18 | 0.36 | 0.0025 |
|  | 7 | 6 | 0.1/0.9 | 2.85 | 0.48 | 0.0025 |
|  | 8 | 6 | 0.1/0.9 | 3.02 | 0.42 | 0.0024 |
| **S3** | 1 | 6 | 0.1/0.9 | 3.39 | 0.17 | 0.0075 |
|  | 2 | 6 | 0.1/0.9 | 3.18 | 0.27 | 0.0051 |
|  | 3 | 6 | 0.1/0.9 | 3.87 | 0.20 | 0 |
|  | 4 | 6 | 0.1/0.9 | 3.48 | 0.18 | 0.0051 |
|  | 5 | 6 | 0.1/0.9 | 4.29 | 0.10 | 0 |
| **S4** | 1 | 6 | 0.1/0.9 | 2.31 | 0.35 | 0.027 |
|  | 2 | 6 | 0.1/0.9 | 1.34 | 0.54 | 0.075 |
|  | 3 | 6 | 0.1/0.9 | 2.35 | 0.40 | 0.018 |
|  | 4 | 6 | 0.1/0.9 | 1.88 | 0.53 | 0.025 |
| **S5** | 1 | 6 | 0.1/0.9 | 1.58 | 0.31 | 0.14 |
|  | 2 | 6 | 0.1/0.9 | 2.39 | 0.21 | 0.056 |
|  | 3 | 4 | 0.1/0.9 | 2.21 | 0.26 | 0.060 |
|  | 4 | 6 | 0.1/0.9 | 1.84 | 0.44 | 0.044 |
|  | 5 | 6 | 0.1/0.9 | 2.56 | 0.35 | 0.015 |
|  | 6 | 4 | 0.1/0.9 | 3.24 | 0.29 | 0.0037 |
| **S6** | 1 | 6 | 0.1/0.9 | 2.58 | 0.17 | 0.052 |
|  | 2 | 6 | 0.1/0.9 | 2.17 | 0.18 | 0.11 |
|  | 3 | 6 | 0.1/0.9 | 2.72 | 0.13 | 0.056 |
|  | 4 | 6 | 0.1/0.9 | 2.32 | 0.18 | 0.079 |
| **S7** | 1 | 8 | 0.1/0.9 | 2.51 | 0.23 | 0.039 |
|  | 2 | 8 | 0.1/0.9 | 2.02 | 0.54 | 0.016 |
|  | 3 | 6 | 0.25/0.75 | 2.35 | 0.40 | 0.018 |
|  | 4 | 6 | 0.25/0.75 | 1.61 | 0.44 | 0.072 |

**Table 3.** Summary of rest sessions conducted.

| Patient | Run # | Duration (mins) | Sampling Rate (Hz) | Instruction |
| --- | --- | --- | --- | --- |
| **S1** | 1 | 5.69 | 1000 | Eyes open+cross fixation |
|  | 2 | 5.43 | 1000 | Eyes open+cross fixation |
|  | 3 | 6.20 | 1000 | Eyes open+cross fixation |
|  | 4 | 8.88 | 1000 | None |
|  | 5 | 7.42 | 1000 | None |
|  | 6 | 7.02 | 1000 | None |
| **S2** | 1 | 5.57 | 1000 | Eyes open+cross fixation |
|  | 2 | 5.61 | 1000 | Eyes open+cross fixation |
|  | 3 | 6.03 | 1000 | Eyes open+cross fixation |
|  | 4 | 7.10 | 1000 | Eyes open+cross fixation |
|  | 5 | 7.76 | 1000 | Eyes open+cross fixation |
|  | 6 | 7.42 | 1000 | None |
|  | 7 | 7.27 | 1000 | None |
|  | 8 | 6.63 | 1000 | None |
| **S6** | 1 | 7.73 | 1000 | Eyes closed |
|  | 2 | 8.67 | 500 | None |
|  | 3 | 8.47 | 500 | None |
|  | 4 | 8.47 | 500 | None |

**Table 4.** Summary of sleep sessions conducted.

| **Patient** | **Run #** | **Duration**<br>**(mins)** | **Sampling**<br>**Rate (Hz)** |
| --- | --- | --- | --- |
| **S1** | 1 | 8.47 | 1000 |
|  | 2 | 8.47 | 1000 |
|  | 3 | 8.48 | 1000 |
|  | 4 | 9.05 | 1000 |
|  | 5 | 8.02 | 1000 |
| **S2** | 1 | 8.83 | 1000 |
|  | 2 | 8.55 | 1000 |
|  | 3 | 4.28 | 1000 |
|  | 4 | 8.18 | 1000 |
|  | 5 | 8.60 | 1000 |
|  | 6 | 5.35 | 1000 |
| **S6** | 1 | 9.10 | 500 |
|  | 2 | 8.22 | 500 |
|  | 3 | 8.50 | 500 |

## References

Andrews-Hanna JR, Reidler JS, Sepulcre J, Poulin R, Buckner RL (2010) Functional-anatomic fractionation of the brain’s default network. Neuron 65: 550–562.

Anticevic A, Cole MW, Murray JD, Corlett PR, Wang XJ, Krystal JH (2012) The role of default network deactivation in cognition and disease. Trends in cognitive sciences 16: 584–592.

Baayen RH, Davidson DJ, Bates DM (2008) Mixed-effects modeling with crossed random effects for subjects and items. Journal of memory and language 59: 390–412.

Biswal B, Yetkin FZ, Haughton VM, Hyde JS (1995) Functional connectivity in the motor cortex of resting human brain using echo-planar MRI. Magnetic resonance in medicine 34: 537–541.

Brainard DH (1997) The Psychophysics Toolbox. Spatial vision 10: 433–436.

Brookes MJ, Woolrich M, Luckhoo H, Price D, Hale JR, Stephenson MC, Barnes GR, Smith SM, Morris PG (2011) Investigating the electrophysiological basis of resting state networks using magnetoencephalography. Proceedings of the National Academy of Sciences of the United States of America 108: 16783–16788.

Buckner RL, Andrews-Hanna JR, Schacter DL (2008) The brain’s default network: anatomy, function, and relevance to disease. Annals of the New York Academy of Sciences 1124: 1–38.

Buckner RL, Krienen FM, Yeo BT (2013) Opportunities and limitations of intrinsic functional connectivity MRI. Nature neuroscience 16: 832–837.

Castellanos FX, Margulies DS, Kelly C, Uddin LQ, Ghaffari M, Kirsch A, Shaw D, Shehzad Z, Di Martino A, Biswal B, Sonuga-Barke EJ, Rotrosen J, Adler LA, Milham MP (2008) Cingulate-precuneus interactions: a new locus of dysfunction in adult attention-deficit/hyperactivity disorder. Biological psychiatry 63: 332–337.

Chai XJ, Castanon AN, Ongur D, Whitfield-Gabrieli S (2012) Anticorrelations in resting state networks without global signal regression. NeuroImage 59: 1420–1428.

Chang C, Glover GH (2009) Effects of model-based physiological noise correction on default mode network anti-correlations and correlations. NeuroImage 47: 1448–1459.

Chen JE, Glover GH, Greicius MD, Chang C (2017) Dissociated patterns of anti-correlations with dorsal and ventral default-mode networks at rest. Human brain mapping 38: 2454–2465.

Christoff K, Gordon AM, Smallwood J, Smith R, Schooler JW (2009) Experience sampling during fMRI reveals default network and executive system contributions to mind wandering. Proceedings of the National Academy of Sciences of the United States of America 106: 8719–8724.

Cole MW, Bassett DS, Power JD, Braver TS, Petersen SE (2014) Intrinsic and task-evoked network architectures of the human brain. Neuron 83: 238–251.

Cole MW, Yarkoni T, Repovs G, Anticevic A, Braver TS (2012) Global connectivity of prefrontal cortex predicts cognitive control and intelligence. The Journal of neuroscience: the official journal of the Society for Neuroscience 32: 8988–8999.

Corbetta M, Shulman GL (2002) Control of goal-directed and stimulus-driven attention in the brain. Nature Reviews Neuroscience 3: 201–215.

Daitch AL, Parvizi J (2018) Spatial and temporal heterogeneity of neural responses in human posteromedial cortex. Proceedings of the National Academy of Sciences of the United States of America 115: 4785–4790.

Dastjerdi M, Foster BL, Nasrullah S, Rauschecker AM, Dougherty RF, Townsend JD, Chang C, Greicius MD, Menon V, Kennedy DP, Parvizi J (2011) Differential electrophysiological response during rest, self-referential, and non-self-referential tasks in human posteromedial cortex. Proceedings of the National Academy of Sciences of the United States of America 108: 3023–3028.

David O, Guillemain I, Saillet S, Reyt S, Deransart C, Segebarth C, Depaulis A (2008) Identifying neural drivers with functional MRI: an electrophysiological validation. PLoS biology 6: 2683–2697.

de Pasquale F, Della Penna S, Snyder AZ, Lewis C, Mantini D, Marzetti L, Belardinelli P, Ciancetta L, Pizzella V, Romani GL, Corbetta M (2010) Temporal dynamics of spontaneous MEG activity in brain networks. Proceedings of the National Academy of Sciences of the United States of America 107: 6040–6045.

Deco G, Jirsa V, McIntosh AR, Sporns O, Kotter R (2009) Key role of coupling, delay, and noise in resting brain fluctuations. Proceedings of the National Academy of Sciences of the United States of America 106: 10302–10307.

Deco G, Jirsa VK, McIntosh AR (2013) Resting brains never rest: computational insights into potential cognitive architectures. Trends in neurosciences 36: 268–274.

Dixon ML, Andrews-Hanna JR, Spreng RN, Irving ZC, Mills C, Girn M, Christoff K (2017) Interactions between the default network and dorsal attention network vary across default subsystems, time, and cognitive states. NeuroImage 147: 632–649.

Downar J, Crawley AP, Mikulis DJ, Davis KD (2000) A multimodal cortical network for the detection of changes in the sensory environment. Nature neuroscience 3: 277–283.

Dykstra AR, Chan AM, Quinn BT, Zepeda R, Keller CJ, Cormier J, Madsen JR, Eskandar EN, Cash SS (2012) Individualized localization and cortical surface-based registration of intracranial electrodes. NeuroImage 59: 3563–3570.

Esterman M, Noonan SK, Rosenberg M, Degutis J (2013) In the zone or zoning out? Tracking behavioral and neural fluctuations during sustained attention. Cerebral cortex 23: 2712–2723.

Fischl B, Sereno MI, Dale AM (1999) Cortical surface-based analysis. II: Inflation, flattening, and a surface-based coordinate system. NeuroImage 9: 195–207.

Fortenbaugh FC, DeGutis J, Germine L, Wilmer JB, Grosso M, Russo K, Esterman M (2015) Sustained Attention Across the Life Span in a Sample of 10,000: Dissociating Ability and Strategy. Psychological science 26: 1497–1510.

Fortenbaugh FC, Rothlein D, McGlinchey R, DeGutis J, Esterman M (2018) Tracking behavioral and neural fluctuations during sustained attention: A robust replication and extension. NeuroImage 171: 148–164.

Foster BL, Rangarajan V, Shirer WR, Parvizi J (2015) Intrinsic and task-dependent coupling of neuronal population activity in human parietal cortex. Neuron 86: 578–590.

Fox MD, Corbetta M, Snyder AZ, Vincent JL, Raichle ME (2006) Spontaneous neuronal activity distinguishes human dorsal and ventral attention systems. ProcNatlAcadSciUSA 103: 10046–10051.

Fox MD, Snyder AZ, Vincent JL, Corbetta M, Van EDC, Raichle ME (2005) The human brain is intrinsically organized into dynamic, anticorrelated functional networks. ProcNatlAcadSciUSA 102: 9673–9678.

Fox MD, Zhang D, Snyder AZ, Raichle ME (2009) The global signal and observed anticorrelated resting state brain networks. Journal of neurophysiology 101: 3270–3283.

Fransson P (2005) Spontaneous low-frequency BOLD signal fluctuations: an fMRI investigation of the resting-state default mode of brain function hypothesis. HumBrain Mapp 26: 15–29.

Fu Z, Wu DJ, Ross I, Chung JM, Mamelak AN, Adolphs R, Rutishauser U (2018) Single-Neuron Correlates of Error Monitoring and Post-Error Adjustments in Human Medial Frontal Cortex. Neuron.

Gratton C, Laumann TO, Nielsen AN, Greene DJ, Gordon EM, Gilmore AW, Nelson SM, Coalson RS, Snyder AZ, Schlaggar BL, Dosenbach NUF, Petersen SE (2018) Functional Brain Networks Are Dominated by Stable Group and Individual Factors, Not Cognitive or Daily Variation. Neuron 98:439–452 e435.

Groppe DM, Bickel S, Dykstra AR, Wang X, Megevand P, Mercier MR, Lado FA, Mehta AD, Honey CJ (2017) iELVis: An open source MATLAB toolbox for localizing and visualizing human intracranial electrode data. Journal of neuroscience methods 281: 40–48.

Hacker CD, Snyder AZ, Pahwa M, Corbetta M, Leuthardt EC (2017) Frequency-specific electrophysiologic correlates of resting state fMRI networks. NeuroImage 149: 446–457.

Ham T, Leff A, de Boissezon X, Joffe A, Sharp DJ (2013) Cognitive control and the salience network: an investigation of error processing and effective connectivity. The Journal of neuroscience: the official journal of the Society for Neuroscience 33: 7091–7098.

He BJ, Snyder AZ, Zempel JM, Smyth MD, Raichle ME (2008) Electrophysiological correlates of the brain’s intrinsic large-scale functional architecture. Proceedings of the National Academy of Sciences of the United States of America 105: 16039–16044.

Hermes D, Miller KJ, Vansteensel MJ, Aarnoutse EJ, Leijten FS, Ramsey NF (2012) Neurophysiologic correlates of fMRI in human motor cortex. Human brain mapping 33: 1689–1699.

Hipp JF, Hawellek DJ, Corbetta M, Siegel M, Engel AK (2012) Large-scale cortical correlation structure of spontaneous oscillatory activity. Nature neuroscience 15: 884–890.

Honey CJ, Newman EL, Schapiro AC (2018) Switching between internal and external modes: A multiscale learning principle. Netw Neurosci 1: 339–356.

Horovitz SG, Braun AR, Carr WS, Picchioni D, Balkin TJ, Fukunaga M, Duyn JH (2009) Decoupling of the brain’s default mode network during deep sleep. Proceedings of the National Academy of Sciences of the United States of America 106: 11376–11381.

Jenkinson M, Beckmann CF, Behrens TE, Woolrich MW, Smith SM (2012) Fsl. NeuroImage 62: 782–790.

Keller CJ, Bickel S, Honey CJ, Groppe DM, Entz L, Craddock RC, Lado FA, Kelly C, Milham M, Mehta AD (2013) Neurophysiological investigation of spontaneous correlated and anticorrelated fluctuations of the BOLD signal. The Journal of neuroscience: the official journal of the Society for Neuroscience 33: 6333–6342.

Keller JB, Hedden T, Thompson TW, Anteraper SA, Gabrieli JD, Whitfield-Gabrieli S (2015) Resting-state anticorrelations between medial and lateral prefrontal cortex: association with working memory, aging, and individual differences. Cortex; a journal devoted to the study of the nervous system and behavior 64: 271–280.

Kelly AM, Uddin LQ, Biswal BB, Castellanos FX, Milham MP (2008) Competition between functional brain networks mediates behavioral variability. NeuroImage 39: 527–537.

Kelly C, Toro R, Di Martino A, Cox CL, Bellec P, Castellanos FX, Milham MP (2012) A convergent functional architecture of the insula emerges across imaging modalities. NeuroImage 61: 1129–1142.

Kiebel SJ, Friston KJ (2004) Statistical parametric mapping for event-related potentials: I. Generic considerations. NeuroImage 22: 492–502.

Kucyi A, Esterman M, Riley CS, Valera EM (2016) Spontaneous default network activity reflects behavioral variability independent of mind-wandering. Proceedings of the National Academy of Sciences of the United States of America 113: 13899–13904.

Kucyi A, Hodaie M, Davis KD (2012) Lateralization in intrinsic functional connectivity of the temporoparietal junction with salience- and attention-related brain networks. Journal of neurophysiology 108: 3382–3392.

Kucyi A, Hove MJ, Esterman M, Hutchison RM, Valera EM (2017) Dynamic Brain Network Correlates of Spontaneous Fluctuations in Attention. Cerebral cortex 27: 1831–1840.

Kucyi A, Schrouff J, Bickel S, Foster BL, Shine JM, Parvizi J (2018a) Intracranial Electrophysiology Reveals Reproducible Intrinsic Functional Connectivity within Human Brain Networks. The Journal of neuroscience: the official journal of the Society for Neuroscience 38: 4230–4242.

Kucyi A, Tambini A, Sadaghiani S, Keilholz S, Cohen JR (2018b) Spontaneous cognitive processes and the behavioral validation of time-varying brain connectivity. Network Neuroscience [in press].

Larson-Prior LJ, Zempel JM, Nolan TS, Prior FW, Snyder AZ, Raichle ME (2009) Cortical network functional connectivity in the descent to sleep. Proceedings of the National Academy of Sciences of the United States of America 106: 4489–4494.

Laumann TO, Snyder AZ, Mitra A, Gordon EM, Gratton C, Adeyemo B, Gilmore AW, Nelson SM, Berg JJ, Greene DJ, McCarthy JE, Tagliazucchi E, Laufs H, Schlaggar BL, Dosenbach NUF, Petersen SE (2017) On the Stability of BOLD fMRI Correlations. Cerebral cortex 27: 4719–4732.

Leonardi N, Van De Ville D (2015) On spurious and real fluctuations of dynamic functional connectivity during rest. NeuroImage 104: 430–436.

Li X, Liang Z, Kleiner M, Lu ZL (2010) RTbox: a device for highly accurate response time measurements. Behavior research methods 42: 212–225.

Liegeois R, Laumann TO, Snyder AZ, Zhou J, Yeo BTT (2017) Interpreting temporal fluctuations in resting-state functional connectivity MRI. NeuroImage.

Logothetis NK, Pauls J, Augath M, Trinath T, Oeltermann A (2001) Neurophysiological investigation of the basis of the fMRI signal. Nature 412: 150–157.

Luke SG (2017) Evaluating significance in linear mixed-effects models in R. Behavior research methods 49: 1494–1502.

Macmillan NA, Creelman DC (2004) Detection Theory: A User’s Guide: Psychology press.

Margulies DS, Ghosh SS, Goulas A, Falkiewicz M, Huntenburg JM, Langs G, Bezgin G, Eickhoff SB, Castellanos FX, Petrides M, Jefferies E, Smallwood J (2016) Situating the default-mode network along a principal gradient of macroscale cortical organization. Proceedings of the National Academy of Sciences of the United States of America 113: 12574–12579.

Maris E, Oostenveld R (2007) Nonparametric statistical testing of EEG- and MEG-data. Journal of neuroscience methods 164: 177–190.

Menon V, Uddin LQ (2010) Saliency, switching, attention and control: a network model of insula function. Brain StructFunct 214: 655–667.

Mesulam MM, Mufson EJ (1982) Insula of the old world monkey. I. Architectonics in the insuloorbito-temporal component of the paralimbic brain. The Journal of comparative neurology 212: 1–22.

Miller KJ, Leuthardt EC, Schalk G, Rao RP, Anderson NR, Moran DW, Miller JW, Ojemann JG (2007) Spectral changes in cortical surface potentials during motor movement. The Journal of neuroscience: the official journal of the Society for Neuroscience 27: 2424–2432.

Mitra A, Kraft A, Wright P, Acland B, Snyder AZ, Rosenthal Z, Czerniewski L, Bauer A, Snyder L, Culver J, Lee JM, Raichle ME (2018) Spontaneous Infra-slow Brain Activity Has Unique Spatiotemporal Dynamics and Laminar Structure. Neuron 98:297–305 e296.

Mukamel R, Gelbard H, Arieli A, Hasson U, Fried I, Malach R (2005) Coupling between neuronal firing, field potentials, and FMRI in human auditory cortex. Science 309: 951–954.

Murphy K, Birn RM, Handwerker DA, Jones TB, Bandettini PA (2009) The impact of global signal regression on resting state correlations: are anti-correlated networks introduced? NeuroImage 44: 893–905.

Murphy K, Fox MD (2017) Towards a consensus regarding global signal regression for resting state functional connectivity MRI. NeuroImage 154: 169–173.

Naidich TP, Kang E, Fatterpekar GM, Delman BN, Gultekin SH, Wolfe D, Ortiz O, Yousry I, Weismann M, Yousry TA (2004) The insula: anatomic study and MR imaging display at 1.5 T. AJNR AmJNeuroradiol 25: 222–232.

Neta M, Miezin FM, Nelson SM, Dubis JW, Dosenbach NU, Schlaggar BL, Petersen SE (2015) Spatial and temporal characteristics of error-related activity in the human brain. The Journal of neuroscience: the official journal of the Society for Neuroscience 35: 253–266.

Nir Y, Fisch L, Mukamel R, Gelbard-Sagiv H, Arieli A, Fried I, Malach R (2007) Coupling between neuronal firing rate, gamma LFP, and BOLD fMRI is related to interneuronal correlations. Current biology: CB 17: 1275–1285.

Nir Y, Mukamel R, Dinstein I, Privman E, Harel M, Fisch L, Gelbard-Sagiv H, Kipervasser S, Andelman F, Neufeld MY, Kramer U, Arieli A, Fried I, Malach R (2008) Interhemispheric correlations of slow spontaneous neuronal fluctuations revealed in human sensory cortex. Nature neuroscience 11: 1100–1108.

Nyberg L, McIntosh AR, Cabeza R, Nilsson LG, Houle S, Habib R, Tulving E (1996) Network analysis of positron emission tomography regional cerebral blood flowdata: Ensemble inhibition during episodic memory retrieval. Journal of Neuroscience 16: 3753–3759.

Oostenveld R, Fries P, Maris E, Schoffelen JM (2011) FieldTrip: Open source software for advanced analysis of MEG, EEG, and invasive electrophysiological data. Computational intelligence and neuroscience 2011: 156869.

Ossandon T, Jerbi K, Vidal JR, Bayle DJ, Henaff MA, Jung J, Minotti L, Bertrand O, Kahane P, Lachaux JP (2011) Transient suppression of broadband gamma power in the default-mode network is correlated with task complexity and subject performance. The Journal of neuroscience: the official journal of the Society for Neuroscience 31: 14521–14530.

Papademetris X, Jackowski MP, Rajeevan N, DiStasio M, Okuda H, Constable RT, Staib LH (2006) BioImage Suite: An integrated medical image analysis suite: An update. The insight journal 2006: 209.

Parvizi J, Kastner S (2018) Promises and limitations of human intracranial electroencephalography. Nature neuroscience 21: 474–483.

Parvizi J, Van Hoesen GW, Buckwalter J, Damasio A (2006) Neural connections of the posteromedial cortex in the macaque. Proceedings of the National Academy of Sciences of the United States of America 103: 1563–1568.

Posner MI, Petersen SE (1990) The attention system of the human brain. Annual review of neuroscience 13: 25–42.

Pruim RH, Mennes M, van Rooij D, Llera A, Buitelaar JK, Beckmann CF (2015) ICA-AROMA: A robust ICA-based strategy for removing motion artifacts from fMRI data. NeuroImage 112: 267–277.

Raccah O, Daitch AL, Kucyi A, Parvizi J (2018) Direct Cortical Recordings Suggest Temporal Order of Task-Evoked Responses in Human Dorsal Attention and Default Networks. The Journal of neuroscience: the official journal of the Society for Neuroscience 38: 10305–10313.

Raichle ME (2015) The brain’s default mode network. Annual review of neuroscience 38: 433–447.

Raichle ME, MacLeod AM, Snyder AZ, Powers WJ, Gusnard DA, Shulman GL (2001) A default mode of brain function. ProcNatlAcadSciUSA 98: 676–682.

Ramot M, Fisch L, Davidesco I, Harel M, Kipervasser S, Andelman F, Neufeld MY, Kramer U, Fried I, Malach R (2013) Emergence of sensory patterns during sleep highlights differential dynamics of REM and non-REM sleep stages. The Journal of neuroscience: the official journal of the Society for Neuroscience 33: 14715–14728.

Ramot M, Fisch L, Harel M, Kipervasser S, Andelman F, Neufeld MY, Kramer U, Fried I, Malach R (2012) A widely distributed spectral signature of task-negative electrocorticography responses revealed during a visuomotor task in the human cortex. The Journal of neuroscience: the official journal of the Society for Neuroscience 32: 10458–10469.

Rosenberg MD, Finn ES, Scheinost D, Papademetris X, Shen X, Constable RT, Chun MM (2016) A neuromarker of sustained attention from whole-brain functional connectivity. Nature neuroscience 19: 165–171.

Rothlein D, DeGutis J, Esterman M (2018) Attentional fluctuations influence the neural fidelity and connectivity of stimulus representations. Journal of cognitive neuroscience 30: 1209–1228.

Schrouff J, Mourao-Miranda J, Phillips C, Parvizi J (2016) Decoding intracranial EEG data with multiple kernel learning method. Journal of neuroscience methods 261: 19–28.

Schrouff J, Rosa MJ, Rondina JM, Marquand AF, Chu C, Ashburner J, Phillips C, Richiardi J, Mourao-Miranda J (2013) PRoNTo: Pattern Recognition for Neuroimaging Toolbox. Neuroinformatics 11: 319–337.

Seeley WW, Menon V, Schatzberg AF, Keller J, Glover GH, Kenna H, Reiss AL, Greicius MD (2007) Dissociable intrinsic connectivity networks for salience processing and executive control. Journal of Neuroscience 27: 2349–2356.

Sestieri C, Corbetta M, Spadone S, Romani GL, Shulman GL (2014) Domain-general signals in the cingulo-opercular network for visuospatial attention and episodic memory. Journal of cognitive neuroscience 26: 551–568.

Shulman GL, Fiez JA, Corbetta M, Buckner RL, Miezin FM, Raichle ME, Petersen SE (1997) Common Blood Flow Changes across Visual Tasks: II. Decreases in Cerebral Cortex. Journal of cognitive neuroscience 9: 648–663.

Sonuga-Barke EJ, Castellanos FX (2007) Spontaneous attentional fluctuations in impaired states and pathological conditions: a neurobiological hypothesis. Neuroscience and biobehavioral reviews 31: 977–986.

Spreng RN, Stevens WD, Viviano JD, Schacter DL (2016) Attenuated anticorrelation between the default and dorsal attention networks with aging: evidence from task and rest. Neurobiology of aging 45: 149–160.

Sripada CS, Kessler D, Angstadt M (2014) Lag in maturation of the brain’s intrinsic functional architecture in attention-deficit/hyperactivity disorder. Proceedings of the National Academy of Sciences of the United States of America 111: 14259–14264.

Thompson GJ, Magnuson ME, Merritt MD, Schwarb H, Pan WJ, McKinley A, Tripp LD, Schumacher EH, Keilholz SD (2013) Short-time windows of correlation between large-scale functional brain networks predict vigilance intraindividually and interindividually. Human brain mapping 34: 3280–3298.

Ture U, Yasargil DC, Al-Mefty O, Yasargil MG (1999) Topographic anatomy of the insular region. Journal of neurosurgery 90: 720–733.

Uddin LQ (2015) Salience processing and insular cortical function and dysfunction. Nature reviews Neuroscience 16: 55–61.

Uddin LQ, Kelly AM, Biswal BB, Castellanos FX, Milham MP (2009) Functional connectivity of default mode network components: correlation, anticorrelation, and causality. Human brain mapping 30: 625–637.

Wang C, Ong JL, Patanaik A, Zhou J, Chee MW (2016) Spontaneous eyelid closures link vigilance fluctuation with fMRI dynamic connectivity states. Proceedings of the National Academy of Sciences of the United States of America 113: 9653–9658.

Weissman DH, Roberts KC, Visscher KM, Woldorff MG (2006) The neural bases of momentary lapses in attention. Nature neuroscience 9: 971–978.

Yeo BT, Krienen FM, Sepulcre J, Sabuncu MR, Lashkari D, Hollinshead M, Roffman JL, Smoller JW, Zollei L, Polimeni JR, Fischl B, Liu H, Buckner RL (2011) The organization of the human cerebral cortex estimated by intrinsic functional connectivity. Journal of neurophysiology 106: 1125–1165.

Zhou Y, Friston KJ, Zeidman P, Chen J, Li S, Razi A (2018) The Hierarchical Organization of the Default, Dorsal Attention and Salience Networks in Adolescents and Young Adults. Cerebral cortex 28: 726–737.

